# The multi-level effect of chlorpyrifos during clownfish metamorphosis

**DOI:** 10.1101/2024.08.02.606305

**Authors:** Mathieu Reynaud, Stefano Vianello, Shu-Hua Lee, Pauline Salis, Kai Wu, Bruno Frederich, David Lecchini, Laurence Besseau, Natacha Roux, Vincent Laudet

## Abstract

Chemical pollution in coastal waters, particularly from agricultural runoff organophosphates, poses a significant threat to marine ecosystems, including coral reefs. Pollutants such as chlorpyrifos (CPF) are widely used in agriculture and have adverse effects on marine life and humans. In this paper, we investigate the impact of CPF on the metamorphosis of a coral reef fish model, the clownfish *Amphiprion ocellaris*, focusing on the disruption of thyroid hormone (TH) signalling pathways. Our findings reveal that by reducing TH levels, CPF exposure impairs the formation of characteristic white bands in clownfish larvae, indicative of metamorphosis progression. Interestingly, TH treatment can rescue these effects, establishing a direct causal link between CPF effect and TH disruption. The body shape changes occurring during metamorphosis are also impacted by CPF exposure, shape changes are less advanced in CPF-treated larvae than in control conditions. Moreover, transcriptomic analysis elucidates CPF’s effects on all components of the TH signalling pathway. Additionally, CPF induces systemic effects on cholesterol and vitamin D metabolism, DNA repair, and immunity, highlighting its broader TH-independent impacts. Pollutants are often overlooked in marine ecosystems, particularly in coral reefs. Developing and enhancing coral reef fish models, such as *Amphiprion ocellaris*, offers a more comprehensive understanding of how chemical pollution affects these ecosystems. This approach provides new insights into the complex mechanisms underlying CPF toxicity during fish metamorphosis, shedding light on the broader impact of environmental pollutants on marine organisms.

**Highlights:**

- Chlorpyrifos (CPF) is an insecticide widely used in agriculture for the past five decades and has adverse effects on marine life and humans
- CPF exposure impairs the formation of characteristic white bands in clownfish larvae, indicative of metamorphosis progression
- During metamorphosis, clownfish larvae lose their elongated body shape and transform into miniature ovoid-shaped adults, these shape changes are less advanced in CPF-treated larvae
- CPF induces systemic effects on cholesterol and vitamin D metabolism, DNA repair, and immunity, highlighting its broader TH-independent impacts

## 1. Introduction

Chemical pollution in coastal waters, stemming from residential, industrial, and agricultural sewage, is a global concern (Tornero & Hanke, 2016; Polidoro et al., 2017; Triassi et al., 2019). Agricultural runoff has been pinpointed as a significant source of pollutants — including of a wide range of pesticides (fungicides, herbicides, insecticides, and antifoulants) — within marine ecosystems (Haynes & Johnson, 2000; Carvalho et al., 2002; Shaw et al., 2010; Bartley et al., 2017; Islam & Tanaka, 2004; Ponce-Vélez & de la Lanza-Espino, 2019; Sabdono et al., 2019), including reefs throughout the globe (Bocquené and Franco, 2005; Kitada et al., 2008, Sheikh et al., 2009; Roche et al., 2011; King et al., 2013; Ponce-Vélez & de la Lanza-Espino, 2019). One of the most extensively used families of insecticides worldwide is the organophosphate family. Despite their efficacy in pest management, organophosphates have also been found to have adverse off-target effects as endocrine disrupting compounds and to lead to mortality, neurotoxicity, and physiological imbalance in both humans and wildlife (Abreu-Villaça & Levin, 2017; Wong et al., 2018; Ubaid Ur Rahman et al., 2021).

Among these endocrine disrupting organophosphates is chlorpyrifos (CPF), an insecticide widely used in agriculture for the past five decades (Juberg et al., 2013a; Kumar et al., 2016; Trasande, 2017). Around 2 million tonnes of CPF are used worldwide annually, 90-99% of which persists in the environment due to low degradability, then leaching into soil, freshwater, and ending up in coastal waters over protracted periods of time (Bosu et al; 2024). Within coral reef ecosystems, comprehensive analyses of pesticides in sediments, water, and biota have revealed the presence of CPF across all compartments, with notably higher occurrences in sediments (Botté et al., 2012; Carvalho et al., 2002; Ponce-Vélez & de la Lanza-Espino, 2019). Occurrences of endocrine disrupting compounds such as CPF in coastal waters could particularly affect coral reef fish whose life cycle includes an oceanic dispersal phase followed by a settled reef phase, during which larvae or young juveniles return to the reef ecosystem (Leis, 2006; Leis et al., 2011). The arrival in the coastal ecosystem is a crucial period involving numerous changes under the control of hormonal processes, known as metamorphosis (Laudet, 2011; McMenamin & Parichy, 2013; Holzer et al., 2017; Besson et al., 2020, Roux et al., 2022).

Metamorphosis is a crucial post-embryonic transition that includes spectacular ecological, morphological, physiological, and behavioural changes (Bishop et al., 2006; Xu et al., 2016; Salis et al., 2018; Campinho, 2019; Nguyen et al., 2022; Roux et al., 2022). Thyroid hormones (THs) are the main hormones triggering metamorphosis in vertebrates, including in coral reef fish (Laudet, 2011; McMenamin & Parichy, 2013, Holzer et al., 2017). For example, during zebrafish metamorphosis, THs promote and regulate skin morphogenesis, adult pigment pattern, visual system, nervous system, skeleton development, and shape craniofacial bones during zebrafish metamorphosis (Baumann et al., 2016; Guillot et al., 2016; Xu et al., 2016; Galindo et al., 2019; Saunders et al., 2019; Aman et al., 2021; Farías-Serratos et al., 2021; Keer et al., 2022). In flatfish, THs control the symmetrical-to-asymmetrical transformation of larvae, including eye migration from one side to the other, and the change from a pelagic to a bottom swimmer (Bao et al., 2011; Xu et al., 2016; Shao et al., 2017). Additionally, functional experiments using compounds impairing THs signalling (called goitrogens) result in limiting cell proliferation, delaying lateral flattening, and decreasing body height during flatfish metamorphosis, demonstrating that THs are necessary for body shape changes (Xu et al., 2016). THs have also been demonstrated to be involved in coral reef fish metamorphosis, such as that of the convict surgeonfish (*Acanthurus triostegus*; Holzer et al., 2017) and of the false clownfish (*Amphiprion ocellaris*; Roux et al., 2023), which they trigger and coordinate at multiple levels.

CPF is mostly known to cause nervous damage in insects through its action as an inhibitor of its main target acetylcholinesterase. Crucially however, it has also been shown to impair the TH signalling pathway in vertebrates (Wołejko et al., 2022; Fortenberry et al., 2012) such as mice (Otênio et al. 2019), zebrafish (Qiao et al., 2021) and convict surgeonfish (Holzer et al., 2017; Besson et al., 2020). To date, the mechanisms of action of CPF on THs signalling is far from being understood, especially given that CPF also affects other hormonal and metabolic pathways (reviewed in Ubaid Ur Rahman et al., 2021). A better understanding of the broad impact of CPF on metamorphosing fish would therefore be needed. Our study thus investigates the impact of CPF at the morphological, hormonal, and transcriptomic level in order to shed light on the intricate interplay between CPF exposure and coral reef fish metamorphosis. To test the effects of CPF we used the anemonefish *Amphiprion ocellaris*, a model organism with well-characterised metamorphic processes (Roux et al., 2019, Roux et al., 2023; Laudet and Ravasi, 2022). We show that CPF effectively impairs metamorphosis progression, exemplified in clownfish by white band formation, and does this by decreasing TH hormone levels. We also show that this effect of CPF can be rescued by TH treatment, showing that this effect on metamorphosis progression is effectively linked to the decrease in TH levels. To gain a better insight on the complexity of CPF effect and its relationship with THs, we further perform transcriptomic analysis after treatment of whole clownfish larvae and show that CPF functions *de facto* as a goitrogen on the TH signalling pathway. Interestingly, we also uncover systemic CPF-specific, TH-independent effects on cholesterol and vitamin D metabolism, as well as signatures of genotoxicity and inflammation, suggesting that CPF simultaneously hits several unrelated targets.

## 2. Materials and Methods

### 2.1. Animal Rearing methods

Experiments were performed on larvae from *A.ocellaris* raised in rearing structures located in Banyuls sur mer (France) and at the LinHai Marine Research Station (Taiwan), maintained as previously detailed in Roux et al, 2021, and as in Roux et al., 2023, with minimal developmental differences observed between the two locations. To minimize these differences, treatments were initiated before metamorphosis, at 5 dph at the Observatoire Océanologique of Banyuls-sur-Mer and 8 dph at the LinHai Marine Research Station. A more detailed description is provided in supplementary Materials and methods (1.1 and 1.2).

### 2.2. Pharmacological treatments, experimental design and sample collection

#### 2.2.1. Chemicals

Larvae were treated with one of the following: 0.1% v/v DMSO (Merck/Sigma-Aldrich, CAT#D5879), MPI (10^-5^M Methimazole, Merck/Supelco CAT#M8506; 10^-6^M Potassium perchlorate KClO_4_, Merck/Sigma-Aldrich CAT#460494; 10^-7^M Iopanoic Acid, Merck/Sigma-Aldrich CAT#14131; in DMSO), T3+IOP (10^-7^M 3,3′,5-Triiodo-L-thyronine, Merck/Sigma-Aldrich CAT#T2877; 10^-7^M Iopanoic Acid, Merck/Sigma-Aldrich CAT#14131; in DMSO), 10-30µg/L CPF (Merck/Sigma-Aldrich CAT#45395; in DMSO), or the combination treatment CPF+T3+IOP. All final concentrations reported above were obtained by dilution of 1000X stocks (i.e. applying stocks at 0.1% v/v). Note that — to simplify the presentation of the results — we refer to the “T3+IOP” and “CPF+T3+IOP” treatments as “T3” and “CPF+T3”, respectively, throughout the main text of the paper (i.e. they both contain 10^-7^M IOP, to suppress background endogenous T3 metabolism).

#### 2.2.2 Experimental design

To investigate the effects of CPF on clownfish metamorphosis and compare it to the artificial reduction of THs signaling, we conducted four different experiments. In the first experiment, three batches of larvae were exposed to increasing concentrations of CPF (10, 20, or 30 µg/L) for 3 and 5 days. These larvae were used to study the effect of CPF on white band formation and TH levels. In our conditions, these 5 days post-treatment (dpt) correspond to the time window of formation of the first 2 white bands in the control condition. In the second experiment, we tested whether T3 treatment could rescue the CPF-induced phenotype by co-exposing pre-metamorphic larvae to CPF alone (20 µg/L), T3 alone (10 mol/L), or a combination of CPF and T3 (20 µg/L and 10 mol/L, respectively) for 5 days. In the third experiment, three batches of larvae were exposed to control conditions, CPF (20 µg/L), MPI (10 mol/L), and T3 alone (10 mol/L). Fish were sampled at 0, 5, 7, 12, and 19 dpt to investigate body shape changes and obtain a complete developmental frame. Finally, the fourth experiment focused on transcriptomic changes in metamorphosing larvae exposed to CPF or pharmacological treatments affecting TH signaling. Larvae were exposed to CPF (20 µg/L), MPI (10 mol/L), or a combination of CPF and T3 (20 µg/L and 10 mol/L, respectively) for 5 days.

#### 2.2.3 Sample collection

On the day of collection, all surviving larvae were collected, immediately euthanised in a petri-dish filled with overdosed tricaine methanesulfonate (MS-222, ethyl 3-aminobenzoate methanesulfonate salt; Merck/Supelco CAT#A5040, 200 mg/L), and quickly imaged (see dedicated section). Each larva was then transferred to a dedicated sterile 2mL microcentrifuge tube filled with 750mL ice-cold Trizol (TRI Reagent; Merck/Sigma-Aldrich CAT#T9424) containing three autoclaved stainless steel beads (EBL Biotechnology CAT#SB2006). Samples were disrupted and homogenised by mechanical agitation in a vibrating bead mill (TissueLyser II, Qiagen CAT#85300, RRID:SCR_018623; 3 min, 30Hz, room temperature). Homogenised sample lysates were stored at - 20°C for THs quantification or at -80°C overnight for RNA extraction. RNA extraction was resumed the following day for 10 selected samples per condition (i.e. 50 samples total: maximum sample number to maintain sufficient per-sample data output (≥ 25 million reads/sample) from a single Illumina P3 flow cell). <u>Selection criteria</u> the 10 larvae actually selected for sequencing were chosen based on homogeneity of end of treatment phenotype. Specifically, for treatments with homogeneous outcomes (DMSO, T3), larvae were selected randomly, while for treatments with heterogeneous outcomes (MPI, CPF, CPF+T3), the most extreme phenotypes were discarded. Specifically, MPI-treated larvae with the most noticeable head-bar were discarded (inferred to be least responsive to MPI). In the CPF treatment, the two larvae looking most- and least-affected were discarded, and all other larvae were sequenced. In the CPF+T3 treatment, larvae looking the least advanced in their metamorphosis were discarded (based on trunk bar development).

### 2.3. Quantification of TH levels

THs were extracted from pools of 5 larvae. Following the protocol developed by Holzer et al, 2017, larvae were weighed, crushed in 500 μl of Methanol with a high-speed benchtop homogeniser (FastPrep 24, MP Biochemicals; RRID:SCR_018599), and centrifuged at 4°C for 10 minutes. Centrifugation was repeated twice, and supernatants from each step were collected and pooled. Then, the pellets were resuspended in a mix of methanol (300μl), chloroform (100 μl) and barbital buffer (150 μl), crushed, and centrifuged at 4°C. Supernatants were collected and preserved with the previous supernatants. Pooled supernatants were then dried at 65°C. Hormones were re-extracted twice from the dried extracts with a mix of methanol, chloroform and barbital buffer, centrifuged. Supernatants were pooled and dried at 65°C. Final extracts were re-suspended in 250 μl of Phosphate Buffer Saline (PBS) and kept at -20°C until quantification. TH concentrations were measured by a medical laboratory of Perpignan (Médipole) using an ELISA kit (Access Free T3, Access Free T4, Beckman Coulter). TH quantification experiments were replicated three times, from three different clutches.

### 2.4. Imaging of treated larvae

Larvae were imaged in a transparent petri dish of overdosed MS-222, just before further processing for RNA extraction (see dedicated section). Pictures were taken on a SC180 digital camera mounted to a SZ61 Zoom Stereomicroscope (RRID:SCR_018950) with a Plan achromatic 0.5x auxiliary objective (110ALK-0.5X-2, CAT#N2165500), transmitted LED illumination, and an external LG-LSLED light source (all Olympus/Evident), using Olympus cellSens Software (RRID:SCR_014551). Scale bars (1mm) were added to all pictures using Fiji/ImageJ (Schindelin et al., 2012; RRID:SCR_002285) Analyze > Tools > Scale Bar. Montages, for each condition, were made using Fiji/ImageJ function Image > Stacks > Stack to Images, on a cropped portion of the image containing the fish.

### 2.5. Morpho-phenotypic markers of metamorphosis

#### 2.5.1 White band formation

White band formation is one of the most salient features of clownfish metamorphosis and has been shown to be regulated by THs (Salis et al., 2018). The adult color pattern of *Amphiprion ocellaris* is characterized by three white bands on an orange body background (Salis et al., 2018, 2019). Using the developmental timing of the white bands formation, specifically at 10 dph under control conditions (DMSO and at the Observatoire Océanologique of Banyuls-sur-Mer), when the fish typically exhibit nearly one full white band, we compared the number of white bands across different experimental conditions with increasing concentrations of CPF (10, 20, or 30 µg/L), T3 (10^-7^M), or MPI (10^-7^M).

#### 2.5.2 Allometric relation and body shape analyses

To assess the impact of treatments related to TH on body morphology, we studied allometric relations and body shape changes. To do so, the standard length (SL, distance from the snout to the middle of the caudal peduncle; mm) of each fish was measured using Fiji/ImageJ from fish photographs. Variation in body shape during metamorphosis was investigated by using landmark-based geometric morphometric methods (James Rohlf & Marcus, 1993). To study allometric body variation, the x and y coordinates of 13 homologous landmarks were digitised using the software TpsDIG2 v2.31 (© 2017, Rohlf). Generalised Procrustes Analysis (GPA) was conducted to align specimens (gpagen function from the R-package geomorph (version 4.0.6); Adams & Otárola-Castillo, 2013; Baken et al., 2021), resulting in a shape dataset. Differences in the pattern of shape variation among treatments were examined through various comparative analyses. Initially, principal component analysis (PCA) was performed on shape variables to explore and visually compare the trajectory of body shape variation among and within treatments along the first two principal component axes. Deformation grids generated by tpsRelw32 V1.70 (© 2017, Rohlf) were used to illustrate and describe shape changes associated with PC axes. Furthermore, we assessed the variation in the pattern of shape transformation among treatments by fitting linear models with all shape variables using geomorph’s procD.lm function, followed by an analysis of variance (ANOVA) conducted with geomorph’s LM.RRPP function (Collyer and Adams, 2018). Subsequently, ontogenetic trajectories defined by days post treatment (0dpt, 5dpt, 7dpt, 12dpt, and 19dpt) in morphospace were compared by using the function trajectory.analysis from geomorph. Pairwise comparisons were then used to assess differences in the amount of body shape changes (i.e. the length of the ontogenetic trajectory in morphospace) among treatments.

### 2.6. Analysis after RNA extraction and sequencing

#### 2.6.1. RNA extraction and sequencing

RNA was extracted and sequenced from A. ocellaris tissues, with full details provided in the Supplementary Materials and Methods (Section 2). The methods are comprehensively described in the Supplementary Materials and Methods, including RNA extraction (Section 2.1), library preparation and sequencing (Section 2.3), and the analysis of bulk RNA-seq data (Section 2.4).

#### 2.6.2. Analysis

Trimmed reads were quantified at the transcript-level using the pseudo-aligner salmon v1.10.1 (RRID:SCR_017036; Patro et al., 2017) against the clownfish *Amphiprion ocellaris* reference transcriptome, using the genome as a decoy (decoy-aware pseudo-alignment; assembly ASM2253959v1; Ryu et al., 2022), as per documentation, and using flags --validateMappings -- seqBias --gcBias. The average mapping rate was 85.12% of total reads.

Salmon output transcript-based quantification files (quant.sf) were imported in RStudio (RRID:SCR_000432; Posit team, 2023) and summarised at the gene level using the tximport function from the tximport package (RRID:SCR_016752; Soneson et al., 2015), referencing the gene models of the *Amphiprion ocellaris* reference genome assembly ASM2253959v1. The counts matrix was obtained by re-calculating counts through the flag countsFromAbundance = "lengthScaledTPM". Gene metadata (names, descriptions) were loaded from a custom-curated reference file based on the integration of Ensembl (still based on assembly AmpOce1.0) and NCBI annotations. This metadata reference is available at the code repository associated with this publication. Counts were processed as a DGEobject (edgeR package, RRID:SCR_012802; Robinson et al., 2010; McCarthy et al., 2012; Chen et al., 2016; Chen et al., 2024) and differences in library size were taken into account by obtaining counts per million (CPM) values (edgeR’s function cpm) to allow a comparable threshold to filter out of lowly expressed genes. Here, we only maintained genes for which at least 10 counts could be detected (arbitrary) in at least 10 samples (our smallest experimental unit being *n*=10 per treatment). This corresponded to a threshold of 1.226291 CPM based on the sample with lowest library size, resulting in a filtering out of 5351 out of 26889 genes (19.9%).

PCA, sample correlation plots, and heatmap visualisations were computed on the filtered count data, taking into account library size differences and correcting against compositional bias through the Median of ratios method internal to the DeSeq2 pipeline (DESeq2 package, RRID:SCR_015687; Love et al., 2014). Counts were then further variance stabilised (DESeq2’s varianceStabilizingTransformation function). PCA and correlation plots were calculated on the most variant genes of the dataset, with a variance threshold set based on the knee of the variance distribution of all genes (kneedle function from kneedle package; Tam <u>etam4260.github.io/kneedle/</u>). Here, this corresponded to a sample variance (s^2^) threshold of 0.21, leading to selecting the top 3418 (15.87%) genes as being “most variant”. <u>Pearson correlation matrix</u>: pairwise Spearman’s rank correlation coefficients between samples were calculated using the “cor” function (base R). <u>PCA plot</u>: PCA was computed on centered, unscaled counts (prcomp function, base R) and the samples’ PC scores were plotted using ggplot2 (RRID:SCR_014601; Wickham 2016), using ggalt’s geom_encircle function (Rudis et al., 2017) to enclose samples from the same treatment within polygons. <u>Heatmaps</u>: all heatmaps were plotted using the function Heatmap from the ComplexHeatmap package (Gu et al., 2016; Gu 2022) reporting z-scores ((counts − mean)/sample standard deviation; base R’s “scale” function with default parameters). <u>Similarity</u>: the similarity categorisation was computed — for each differentially expressed gene — by calculating the average z-score value under each treatment, and calculating z-score differences between treatments (i.e. distances in standard deviation units). If the closest value was more than one standard deviation away, the treatment response was categorised as unique to that treatment (similar to neither). If the values in both other treatments were within 1 standard deviation from the one of the treatment of interest, the treatment response was not adjudicated to either (similar to both). Otherwise, similarity was adjudicated to the treatment with the closest distance.

Differential Expression analysis was computed on the filtered count data, normalised using the trimmed mean of M values (TMM) method with reference to the sample with smallest library size to remove compositional bias (Robinson, Oshlack, 2010; through edgeR’s calcNormFactors function), and then processed through the limma-voom pipeline (voom function from limma package, RRID:SCR_010943; Ritchie et al., 2015) feeding the model matrix −1+treatment given that no dominant batch effect was apparent by PCA exploratory analysis (i.e. no effect stronger than the treatment effect). <u>Differential Expression</u>: Linear regression models were fit to the counts data, using voom’s mean-variance precision weights (limma’s lmFit). Pairwise treatment−DMSO contrast matrices were built (limma’s makeContrasts), and computed coefficients (limma’s contrasts.fit) were moderated by Empirical Bayes smoothing of standard errors (limma’s eBayes). A gene was categorised as differentially expressed if showing a log_2_FC < −1 or > +1 (i.e. two folds or more), and associated with a Benjamini-Hochberg-corrected (adjusted) p value < 0.05 (limma’s topTable adjust.method = "BH"). Gene-enrichment and over-representation analysis: was performed based on the Ensembl "biological_process" Gene Ontology annotation for *Amphiprion ocellaris* (“aocellaris_gene_ensembl” dataset; useMart and getBM functions from biomaRt package, RRID:SCR_019214; Durinck et al., 2005; Durinck et al., 2009). Given that Ensembl annotations are based on a different (earlier) genome assembly (AmpOce1.0), genes with no matching Ensembl ID could not be taken into account. Enrichment analysis was performed using the enricher function from the clusterProfiler package (RRID:SCR_016884; Yu et al., 2012; Wu et al., 2021) with default parameters (minGSSize = 10, maxGSSize = 500, universe = all Ensembl genes) and qvalueCutoff = 0.05. Highlighted pathways were consistent across alternative enrichment and over-representation tests (see notebook). Each set of differentially-expressed genes was also parsed and annotated manually to complement GO-based approaches. <u>Venn plots</u> were plotted with the ggVennDiagram function from the ggVennDiagram package (Gao et al., 2021), or venneuler from the venneuler package (preserving relative scale).<u> Alluvial plots</u>: were plotted using functions from the ggalluvial package (Brunson, Read 2023; Brunson 2020). Of genes with a statistically-significant change in expression in both of the treatments compared, expression was classified as “increased” if the change in expression (compared to control) in the second treatment was stronger than the one in the first treatment (higher for genes with positive change, or lower for genes with negative change), “attenuated” if the change in expression was weaker, and “reversed” if the change in expression was of opposite sign across the two treatments.

## 3. Results

### 3.1. CPF impairs white band formation in clownfish in a TH-dependent manner

To determine CPF effects during metamorphosis in clownfish, we exposed larvae for 5 days (from 5dph, i.e. before metamorphosis), to different concentrations of CPF (10µg/L, 20µg/L or 30µg/L). We observed that fish exposed to CPF show a delay in the appearance of white bands when compared to DMSO vehicle controls (**Fig. 1A-D**). This effect is dose-dependent with 100% of the fish exhibiting one or two bands in controls, 60% at 10µg/L, 52% at 20µg/L, and 25% at 30µg/L of CPF (**Fig. 1E**). Fish in the control conditions have significantly more white bands than fish exposed to CPF, at all pesticide concentrations (Control vs CPF-10µg/L, p-value = 0.0007559; Control vs CPF-20µg/L, p-value = 0.0006033; Control vs CPF-10µg/L, p-value = 5.29E-06; χ2 test). These results demonstrate that CPF alters the timing formation of white bands during clownfish metamorphosis. We indeed observed that white bands eventually appear, but at much later time points (after 19 days of treatment, **Fig. S1**).

**Figure 1.**
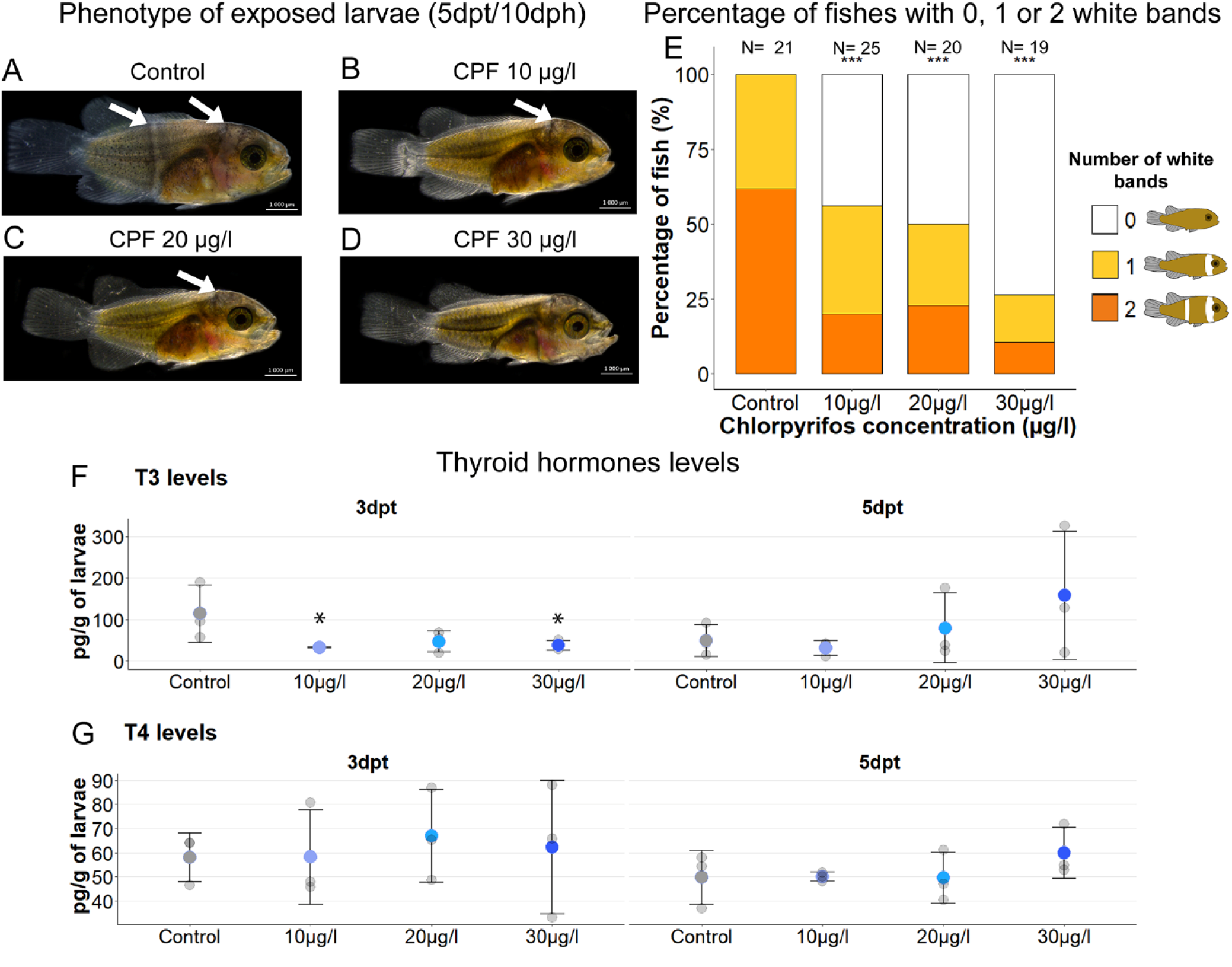
Chlorpyrifos treatment leads to a delay in white band formation and induces TH levels disruption. (A-D) Stereomicroscope images of larvae at 5 dpt in control (A) or CPF at 10µg/L (B), 20µg/L (C), and 30µg/L (D). White arrows indicate the presence of white bands. Scale bar = 1000µm. (E) Percentage of larvae having 0 (white), 1 (yellow) or 2 (orange) white bands. (nDMSO=21, nCPF 10µg/l =25, nCPF 20µg/l=22, nCPF 30µg/l=19 individuals). Chi2 tests are significant between DMSO and CPF 10µg/L, CPF 20µg/L or CPF 30µg/L (p-value <0.001). Significant differences are indicated by a star (***; p-value<0.001). (F-G) TH levels (F) T3, (G) T4, at 3 and 5dpt. Data are indicated as mean (coloured circles) ± SD (error bars), and grey circles indicate each data point. Significant differences are indicated by a star (*, p-value <0.5, Dunn tests, Holm method).

As white band formation has been shown to be dependent on THs (Salis et al., 2021) we next evaluated whether CPF was effectively able to decrease TH levels during clownfish metamorphosis. Accordingly, we again exposed pre-metamorphosis larvae to increasing concentrations of CPF (10, 20 or 30µg/L CPF) and we quantified T3 and T4 levels at 3dpt and 5dpt. At 3dpt, T3 levels were significantly reduced in larvae exposed to 10 and 30µg/L of CPF compared to the control (**Fig. 1F**, P-value <0.05), and they were reduced, albeit non significantly, after treatment at 20µg/L CPF. T4 3dpt levels remained instead stable for all CPF concentrations (**Fig. 1G**, P-value >0.05). At the end of the exposure period (5dpt), no significant differences in T3 and T4 were observed between the control and CPF conditions. These results, showing a transient effect on T3 but not on T4, are strikingly similar to those observed in surgeonfish larvae (Holzer et al., 2017).

Having observed a concomitant effect of CPF on white band formation and T3 levels, we next tested whether a causal relationship exists between the two. We therefore tested whether T3 treatment was able to rescue the CPF-induced phenotype. For this, we conducted co-exposure treatment of pre-metamorphic larvae (at 5dph) with CPF alone (20µg/L), T3 alone (10^-7^mol/L) and a mix of CPF + T3 (20µg/L, 10^-7^mol/L, respectively) for 5 days. Again, the effect on white bands caused by CPF was significant compared to the control (**Fig. 2**; P-value: 0.005) and T3 treatment accelerated white band formation as observed in Salis et al., 2021. Interestingly, co-exposure of T3 and CPF rescued the phenotype induced by CPF alone: co-treated larvae are similar to the control group (**Fig. 2**; P-value: 0.855). This result clearly shows that the CPF effect on white band formation is causally linked to the observed decrease of T3.

**Figure 2.**
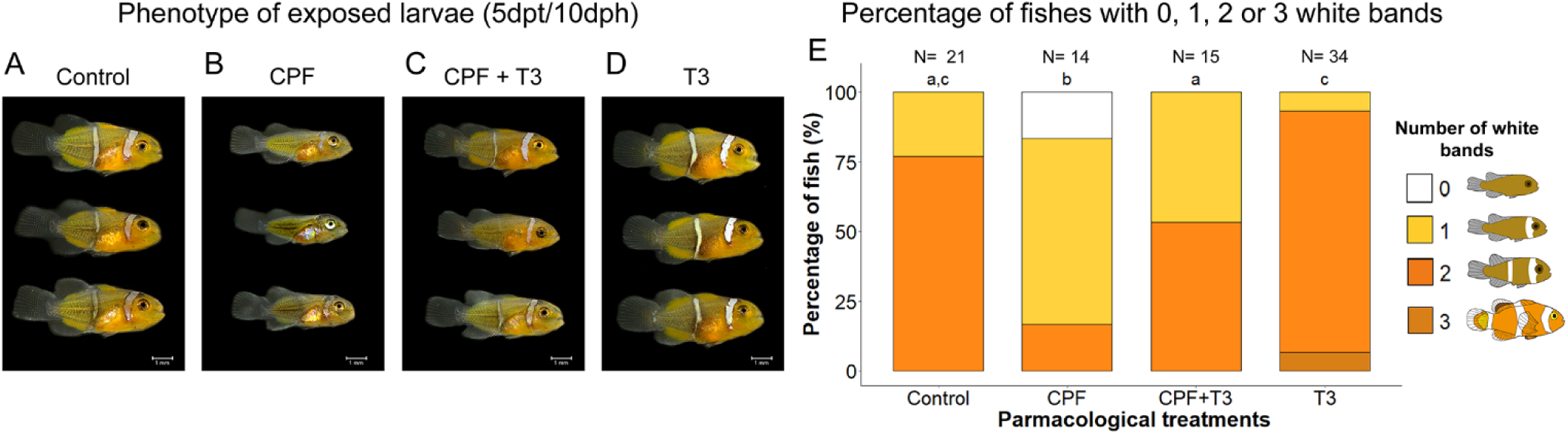
Additional T3 treatment rescues the white band phenotype disrupted by Chlorpyrifos treatment. (A-D) Stereomicroscope images of larvae at 5 dpt in control (A), (B) CPF 20µg/L, (C) CPF 20µg/L and T3 10^-7^mol/L or (D) T3 at 10^-7^mol/L. (E) Percentage of larvae having 0 (yellow), 1 (blue) or 2 (orange) white bands. (nDMSO=21, nCPF 20µg/L =14, nCPF + T3 =15, nT3 =34 individuals). Letters refer to significant differences among treatments calculated by a chi² test.

### 3.2. CPF alters the pattern of ontogenetic shape changes in clownfish larvae

Although white band formation is the most noticeable phenotypic effect of THs in clownfish, it is not the only readout of metamorphosis. We therefore explored the effect of CPF exposure on the known changes in larval body shape that occur during clownfish metamorphosis (Roux et al., 2023). To this aim, we used a geometric morphology approach based on 13 anatomical landmarks (**Fig. S2**) comparing larvae exposed to CPF (20µg/L) to control larvae treated with DMSO, as well as to larvae treated with T3 (10^-7^mol/L) or with a cocktail of goitrogens that impair TH production (Methimazole, Potassium Perchlorate, Iopanoic Acid: MPI, 10^-7^mol/L), as described in Roux et al., 2023. Larvae were treated at 8 dph, for 19 days (0dpt to 19dpt) to fully capture the temporality of the developmental trajectory affected by CPF.

Developmental trajectories are clearly discernible in the shape space defined by the two first principal components (**Fig. 3A**). The main shape changes revealed by PC1 (45.3 % of total shape variation), clearly associated with fish development, concerned modification of relative body depth (**Fig. 3D-E**). PC2 (9.15% of total shape variation) captured changes associated with head shape (**Fig. 3B-C**). Samples from Control treatment draw a clear developmental trajectory along PC1 from 0dpt to 19dpt. Small variation along PC2 is also observed between 5dpt and 7dpt, revealing transient head shape transformation during clownfish metamorphosis. Globally, during metamorphosis, clownfish larvae lose their elongated body shape and transform into miniature ovoid-shaped adults. In CPF-exposed larvae, the separation of sampling periods along PC1 is far less pronounced than in control. Notably, 5dpt individuals remain mixed with 0dpt, 7dpt, and 12dpt fish, suggesting a profound alteration of the pattern of shape changes. Overall, shape changes are less advanced in CPF-treated fish than in control. Similarly, fish exposed to MPI are less advanced than the control condition, and less regularly distributed along PC1 as a function of age. Larvae exposed to T3 are, in contrast, clearly distributed according to age along the PC1 axis. In particular, 5dpt larvae are totally separated from 0dpt ones, and only marginally mixed with 7dpt fish, reflecting an acceleration of the shape changes in this treatment. Moreover, we noticed that the PC2 values of T3-treated fish have a stronger increase at 5dpt compared to the control, an effect we do not see in CPF or MPI-treated fish.

**Figure 3.**
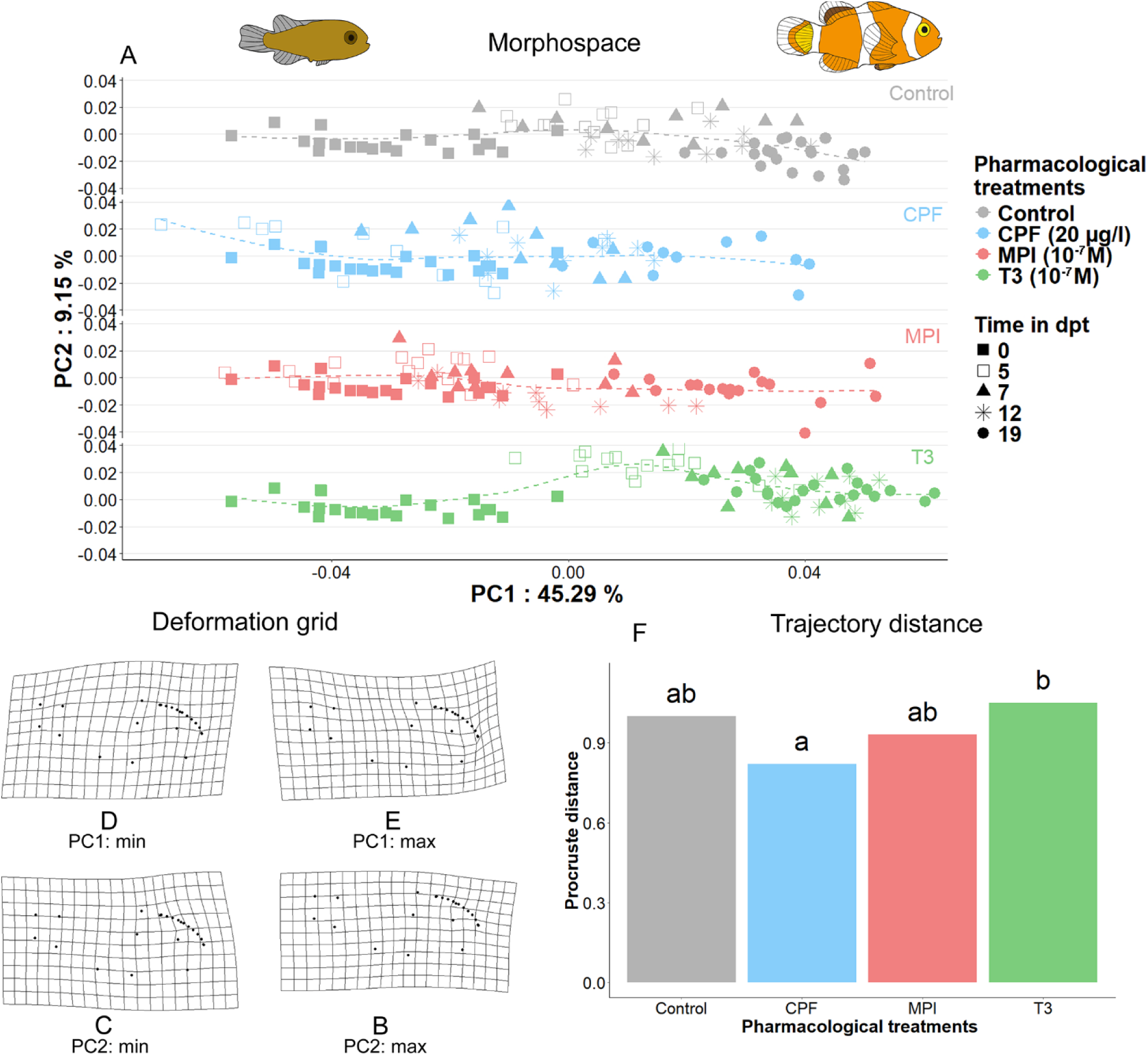
CPF alters body shape changes associated with metamorphosis. (A) plots of the first two principal components (PC1 and PC2) defining the shape space. Larvae of each treatment are separated into different scatter plots (grey = Control DMSO, light blue =CPF (20µg/l), red = MPI (10^-7^mol/L)(Methymazole+Potassium Perchlorate+Iopanoic acid, green = T3 (10^-7^mol/L). The shape of the point illustrates the time in dpt (filled squares: 0dpt, empty squares: 5dpt, triangles: 7dpt, stars: 12dpt, circles: 19dpt). The percentage of shape variation explained by each PC is provided on the y and x-axes of the plot. Deformation grids illustrate shape changes associated with each PC: (B) PC2 max, (C) PC2 min, (D) PC1 min, (E) PC1 max. (F) Length of the ontogenetic trajectories translating the amount of shape variation observed after 19dpt. Letters show significant differences among treatments.

Multivariate linear models reinforce our visual exploration of morphospace occupation. The models confirm that body shape varies across time and treatment (**Table S1**). Treatment significantly affects the rate of shape changes during the studied period of development (F= 10.40, Z=8.41 p-value=0.001; **Table S1**). Developmental trajectories within the morphospace were delineated by connecting four linear segments representing the five sampling periods (0dpt, 5dpt, 7dpt, 12dpt, and 19dpt). Comparative analysis of these time-segmented developmental trajectories revealed differences in terms of distance (**Fig. 3F**). Larvae exposed to CPF exhibited the shortest trajectory (0.055), followed by MPI larvae (0.062), indicating that the amount of shape variation observed at 19dpt is lower than the shape transformation observed under the control condition during the same period of time (0.067). The longest trajectory was observed in T3-treated larvae (0.070). Significant differences were observed between CPF and T3 treatment (d= 0.0148, Z= 1.7385, p-value = 0.033). Taken together, these results highlight the similarity of CPF and MPI treatment on shape transformation, reinforcing the notion that CPF acts here too by decreasing T3 levels.

### 3.3. CPF triggers unique transcriptomic changes in metamorphosing larvae, with mixed features of both TH-activation and TH-blockade

The effects described above, highlight a strong similarity between CPF- and MPI treatments (decreased TH levels, impaired metamorphosis). Still, we do not expect these readouts to capture the full complexity of the mode of action of the pesticide. Accordingly, we performed bulk transcriptomic analysis of whole CPF-, T3-, and MPI-treated larvae, as well as CPF+T3 (rescued treatment; same concentrations as used throughout, see materials and methods). These analyses are likely able to provide a more global view of the effects of CPF on clownfish larvae and on the metamorphic process. An overview of the main features of the bulk RNAseq dataset is presented in **Fig. 4**, and the phenotype of treated larvae (similar to those observed in the corresponding treatments in Roux et al., 2023) is shown in **Fig. S3**. Summary data relating to the technical aspects of library preparation (library size, mapping rates, etc.) is provided in the materials and methods, and in the documents associated with this publication.

**Figure 4:**
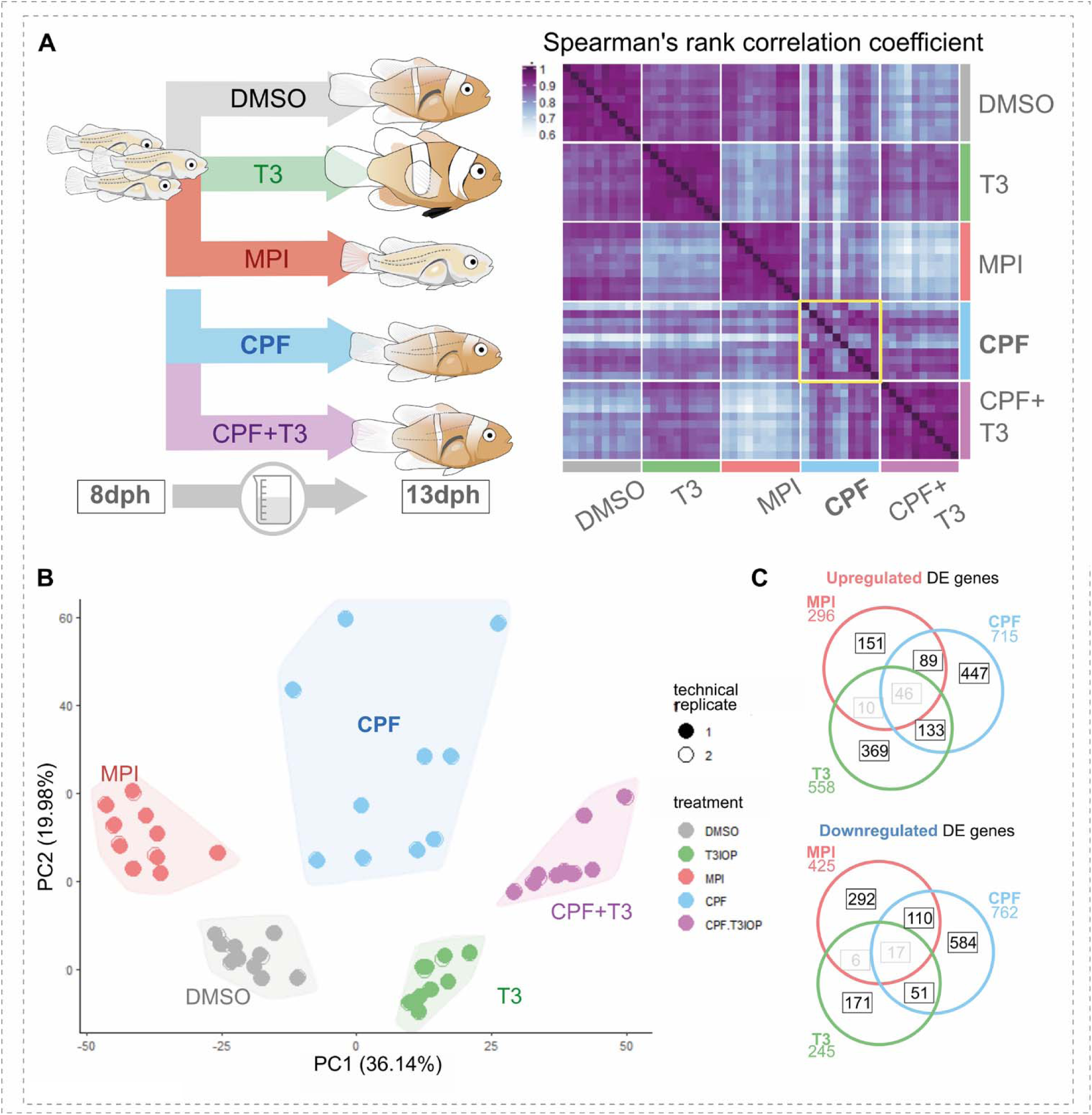
CPF triggers unique transcriptomic changes in metamorphosing larvae, with mixed features of both TH-activation and TH-blockade. **A) left:** schematic of the experimental treatment of samples processed for bulkRNAseq analysis; **right**: correlation matrix showing pairwise Spearman’s rank correlation coefficients, calculated on the top most variable genes across all conditions. Area framed in yellow: correlation coefficient values among CPF-treated larvae. **B)** PCA plot showing the distribution of samples across PC1 and PC2, based on the top most variable genes across all conditions. Technical replicates: data from the two lanes of the same chip. **C)** Venn diagrams of the number of differentially expressed (DE) genes under each treatment (compared to vehicle controls), and their overlaps between treatments. Genes that changed in the same direction in treatments with expected opposite effects (MPI and T3) are shaded out.

We observe that, unlike T3 and MPI treatment, exposure to 20µg/L of CPF results in a rather heterogeneous transcriptomic response in clownfish juveniles. Indeed, the transcriptomes of CPF-treated larvae show the most variable within-treatment correlation coefficients, as well as amongst the lowest ones (section framed in yellow in **Fig. 4A**). Such a high heterogeneity in the response to CPF treatment is similarly reflected by the wider spread of CPF samples across the main principal components plane (**Fig. 4B**), though each treatment condition clusters distinctly. To our surprise, we also notice that, within such heterogeneity, transcriptomic changes resulting from CPF exposure not only correlate to MPI but also to T3, and to similar degrees (**Fig. 4A**). In summary, even though the pesticide clearly results in an impaired-metamorphosis phenotype at the phenotypic level (see previously, **Fig. 1** and **Fig. S3**), the effects of the pesticide may not be correctly described as being MPI-like or TH-like at the transcriptome level. Rather, CPF appears to induce unique transcriptomic changes involving — among others — TH-sensitive genes (i.e. genes also modulated by T3 or MPI).

While T3 has mostly an activating action on gene expression (70% of differentially expressed (DE) genes are upregulated) and MPI a repressive action on gene expression (59% of DE genes are downregulated), CPF is seen to have a more mixed action (48% DE genes upregulated, 52% downregulated) (**Fig. 4C**). Notably, it shares almost equal proportions of DE genes with either of the two opposite treatments (16.7% and 17.7% of its DE shared with T3 and MPI DE genes, respectively), suggesting that CPF acts mimicking both MPI, and paradoxically T3 (**Fig. 4C**). Accordingly, the pattern of expression of differentially expressed genes under CPF treatment is mostly unique to the CPF condition. Yet expression levels can at times be seen to be similar to those of T3-treated samples, at times instead to those in MPI-treated samples. In this, CPF thus correlates to the effects of both treatments, though globally remaining distinct (**Fig. S4**).

Overall, we thus find that CPF triggers a distinct transcriptomic response in metamorphosing clownfish larvae (which respond heterogeneously to the pesticide), and though MPI-like effects may be seen at the phenotypic level, at the transcriptome level these also involve responses normally caused by increased TH levels.

### 3.4. CPF has goitrogenic effects on the larval HPT axis, though also shares systemic effects with T3

Given the role played by TH in driving vertebrate metamorphosis, a role we have previously extensively documented in clownfish (Roux et al., 2023), and knowing the link between CPF and TH action in a number of experimental settings (Holzer et al, 2017; Besson et al, 2020), we first sought to fully characterise the points of overlap between the genes affected by CPF treatment, and those regulated by TH or MPI. That is, within the unique transcriptomic signature induced by CPF, we isolate the most likely candidates that could explain the effects of CPF on metamorphosis progression. For this, we focused on genes involved in the TH-signalling pathway itself, as well as on the genes responsive to this pathway, such as iridophore genes (Salis et al., 2019; Libin et al., 2022), and on the genes involved in the upstream control of TH production (Hypothalamo-Pituitary-Thyroid axis, HPT; see Blanton and Specker, 2007). In vertebrates, particularly in amphibians, a connection also exists between THs and the ACTH/corticoid (Hypothalamo-Pituitary-Interrenal axis, HPI) axes. Indeed, the CRH (the hypothalamic peptide controlling the HPI axis) has also been found to control TSH and TH production (Larsen et al., 1998; De Groef et al., 2006; de Jesus 1990) (**Fig. 5A**). Though the presence or activity of this link in fish is still debated (Huerliman et al. 2024), we nonetheless analysed the effect of CPF on both axes.

**Figure 5:**
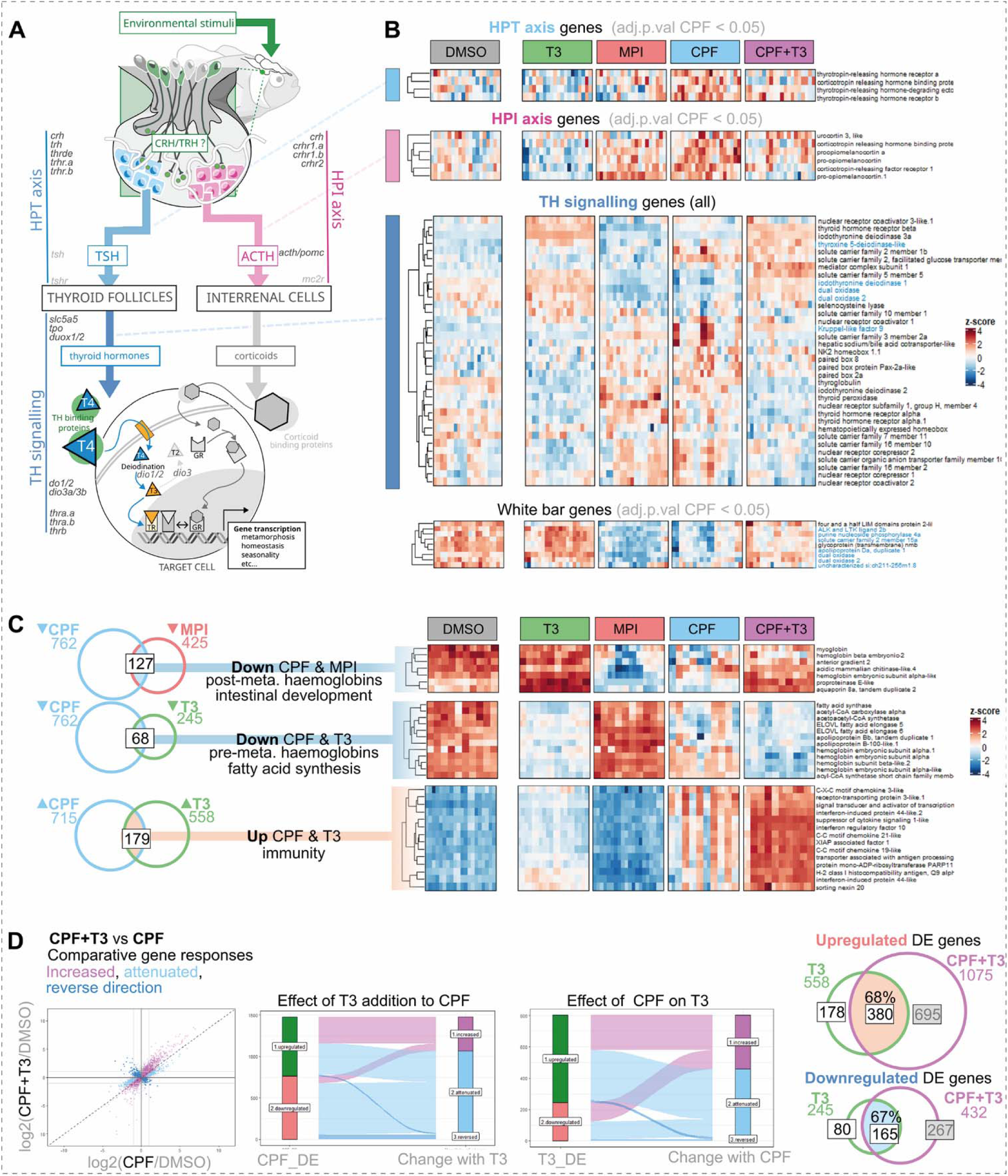
CPF has goitrogenic effects on the larval HPT, though also shares systemic effects with T3. **A)** schematic of the main components of the HPT (light blue) and HPI (magenta) axis in clownfish, and of the TH signalling pathway (dark blue). The role of CRH as an activator of the HPT axis is still debated in teleosts. Genes not detected in the dataset are shaded light-grey **B)** expression pattern (z-scores) of genes involved in TH signalling (all members of the pathway), and in white band formation, HPT axis, and HPI axis (only genes showing significant change in CPF with respect to vehicle controls. Genes highlighted in blue have abs(logFC)>1 (i.e. they pass threshold of differential expression) **C) Left**: Venn diagrams showing the overlap (to scale) between genes differentially expressed in CPF and in either T3 or MPI. **Right**: expression pattern (z-scores) of selected metamorphosis-related genes at different treatment intersections. **D) Left**: scatterplot comparing the magnitude and direction of change of each gene to CPF, and to CPF+T3. Genes changing in opposite directions to the two treatments are indicated in dark blue (“reverse direction”). For genes with coherent direction of change, genes whose response is magnified by the addition of T3 are indicated in purple (“increased”, above the diagonal), genes whose response is dampened by the addition of T3 are indicated in light blue (“attenuated”, below the diagonal). **Middle**: alluvial plots linking DE genes upon CPF treatment, to their response when T3 is also present (i.e. rescue treatment); and vice versa. **Right**: Venn diagrams showing the overlap (to scale) between the genes changing under T3 treatment, green, and those changing when T3 is supplied on a CPF background.

At the level of the expression of TH-signalling pathway components we critically find that CPF mimics the effects of goitrogens (MPI), especially in the downregulation of TH synthesis genes (**Fig. 5B, niddle panel**). Specifically, the observed downregulation of the iodothyronine deiodinase *dio1*, of the two dual oxidases *duox1* and *duox2*, as well as the decrease in expression of the sodium-iodide symporter *slc5a5* suggests that CPF, like MPI, impairs T3 production, as observed in **Fig. 1**. Consistently, we observe a significant increase of expression of embryonic development genes normally associated with thyroid growth and differentiation (*nkx2.1, pax8, pax2a*), likely reflecting compensatory changes to hypothyroidism. Similar changes have been recently described in zebrafish in response to 2-ethylhexyl diphenyl phosphate (EHDPP) another organophosphate pesticide (Shu et al., 2024). Furthermore, we see that the goitrogenic effect of CPF can be completely rescued by T3 supplementation: CPF+T3 samples show a reversed TH pathway signature analogous to that of T3-only samples (**Fig. 5B, middle panel**). This latter observation therefore suggests that CPF does not directly damage thyroid follicular cells, but rather only impairs their ability to produce active THs.

Interestingly, these data are also sufficient to explain the effect of CPF treatment (and T3 rescue) on white band formation described in previous sections (**Fig. 2**). Indeed, *duox1*, a gene found to mediate band patterning dynamics during clownfish metamorphosis (Salis et al., 2019), and which is a known regulator of iridophore development in zebrafish (Chopra et al., 2019; Salis et al., 2019), is downregulated by CPF in a T3-dependent fashion. In addition, we observe that all iridophore development genes downregulated by MPI, and therefore correlated to the observed failure of white band development (*fhl1b, alkal2b, pnp4a, slc2a15a, gpnmb, apod1a, saiyan/si:ch211-256m1.8*), are similarly downregulated by CPF, again in a T3-dependent manner (**Fig. 5B, bottom panel**). The relative level of expression of these genes within the CPF condition qualitatively correlates with the actual band phenotype of the larvae themselves, since the larvae with the lowest level of expression are also the ones with the faintest bands (larvae 1,5,6; highlighted in yellow in **Fig. S3**).

It is less clear whether the effect of CPF on TH production derives from upstream disruption of the HPT or HPI axes. Indeed, of the hypothalamic peptides controlling TSH production, TRH and CRH expression levels do not have significant change compared to control. A significant CPF-treatment-specific increase in expression, though heterogeneous and of low magnitude, is however detected for the receptors of TRH and CRH on pituitary cells, and for the TRH-degrading ectoenzyme (**Fig. 5B, top panels**). Still, the expression of TSH itself is undetectable, as well as that of the receptor that would transduce this signal to thyroid follicles (*thsr*). Accordingly, and though we see clear goitrogenic effects at the level of the TH-synthesis and signalling pathway, we believe that our data is unable to address whether such effect may (also) derive from disruption of the hypothalamic/pituitary control of TH-production.

Having found that CPF effectively acts as a goitrogen on the larval TH-pathway genes, we note that comparison between differentially expressed gene sets across treatments had shown that CPF only partially recapitulates the transcriptomic signature of the MPI goitrogen mix: that is, only 36% of MPI regulated genes are also regulated by CPF (**Fig. 4C**). Based on the available clownfish Gene Ontology reference, no pathway appears to be significantly enriched within this conserved transcriptional response (not shown; see code associated with this publication). Literature-based manual imputation of function (for the genes where this was possible at the time of writing) highlights disparate functions that, to the best of our knowledge about teleost metamorphosis, we are unable to summarise or to link to specific modes of action of MPI. We do however identify genes with clear developmental or metamorphic roles, as also based on our previous work on the topic (Roux et al., 2023; Salis et al., 2019). Accordingly, we highlight that amongst CPF’s MPI-like effects — in addition to those on white band pigmentation mentioned above — are effects indicating an impaired switch to post-metamorphosis haemoglobins (also discussed later), as well as downregulation of genes involved in intestinal development (anterior gradient 2 *agr2*), intestinal/digestive function (aquaporin 8a *aqp8.2*; acidic mammalian chitinase 3/4-like; two trypsinogen1-like; elastase 2a-like), and ion regulation (sodium-phosphate symporter *slc34a2b*) (**Fig. 5C**). A complete list of all genes in each treatment intersection is provided as a **Supplementary File**.

The observation that CPF acts as a goitrogen on the larval TH-signalling pathway yet sharing transcriptomic effects with T3 treatment (i.e. TH activation; 30% of T3 upregulated and downregulated genes combined) may appear paradoxical. Yet this suggests that some of the transcriptional effects observed in the T3 condition can be triggered by T3-independent processes (i.e. are not necessarily caused by increased TH levels). Again, we focus on genes we know to be intimately associated with clownfish metamorphosis, or with clear function during clownfish development. As shown in **Fig. 5C**, both CPF and T3 lead to downregulation of lipid metabolism genes (specifically, the fatty acid synthase gene *fasn*, and the fatty acid elongases *elovl5* and *elovl6* mediating short- and long-chain unsaturated fatty acid synthesis, but also of the lipid precursor generating enzymes acetoacetyl-CoA synthetase and acetyl-CoA carboxylase). We have previously described how the T3-regulated natural metamorphosis of clownfish larvae is characterised by a major metabolic shift in energy production involving divestment from lipogenesis/fatty acid elongation (Roux et al., 2023). This is what we also recover here. Accordingly, the apparent incongruence between the CPF-treated larvae and a T3-like transcriptomic responses simply appear to represent the overlap between — on one hand — the natural switching-off of specific metabolic functions (dependent on an active TH signalling pathway) and — on the other hand — CPF-specific transcriptional response (TH signalling independent). The shared upregulation of immune genes (**Fig. 5C**) may similarly reflect the overlap between the T3-regulated natural development of immunity during metamorphosis, and the damage-induced upregulation of these components in larvae exposed to CPF.

Another T3-like effect of CPF, yet decoupled from accompanying systemic changes that would however happen during TH-regulated metamorphosis, is seen in its downregulation of pre-metamorphosis haemoglobins. Fish metamorphosis is characterised by a switch of the haemoglobin gene sets expressed, endowing the larva with a specialised post-metamorphosis (haemo)globin complement that, in clownfish, has been suggested to be tailored to the hypoxic environment of coral reefs (Downie et al., 2023). Here, though CPF leads to the downregulation of pre-metamorphosis globins just like T3, it also leads, unlike T3, to the downregulation of post-metamorphosis ones (as noted in the MPI-like effects; **Fig. 5C**). Just as what we see for lipogenesis genes, it again appears that at least some of the (unexpected) similarities between T3 and CPF treatments may stem from the partial overlap between i) the switching-off phase of T3-induced physiological transitions, and ii) the widespread switching off of gene functions under CPF. Of interest, comparing the transcriptional response of CPF vs CPF+T3 (and vice versa) shows that exogenous T3 carries a strong attenuating power on the transcriptional responses induced by CPF (**Fig. 5D, left**), especially on the genes downregulated by the pesticide. Still, and though T3 supplementation to CPF can recover a T3-like TH signalling pathway status (**Fig. 5B**), it does not fully recapitulate the T3-only status (**Fig. 5D, right**), and only recovers around 68% of the differentially expressed genes of the T3 condition.

In conclusion, we find that CPF acts as a goitrogen on the TH signalling pathway of clownfish larvae, impairing normal T3-mediated processes such a white band formation, intestinal development, and blood oxygen transport, in an analogous fashion to the reference compound mix MPI. The goitrogenic effect on the larval TH signalling pathway can be fully rescued by the addition of exogenous T3: all pathway components are expressed at T3-like levels even in the presence of CPF. Still, T3 addition and the recovery of an active TH-signalling status is not sufficient to contrast all CPF-induced responses. Conversely, we also find that while CPF may appear to recapitulate T3-regulated changes that normally accompany metamorphosis (e.g. lipogenesis switch off, haemoglobin switch), in contrast to T3 effects, this clearly occurs in a way that is not coordinated at the organism level. Still, it is evident that CPF triggers effects that cannot be attributable to TH signalling pathway modulation as they are unaffected by either T3 or MPI (**Fig. 5C**). In fact, these are the majority of the differentially expressed genes detected in our study.

### 3.5. CPF affects cholesterol and vitamin D synthesis pathways, DNA repair, and immunity

Regardless of the effect of CPF on the larval TH signalling pathway and TH levels, GO-driven analysis of differentially expressed genes between CPF-treated and DMSO-control juveniles highlights three major effects of the organophosphate pesticide, which in fact dominate the transcriptional response of the larvae. These involve: i) cholesterol and vitamin D synthesis, ii) DNA damage/repair, and iii) immunity/antiviral responses (**Fig. 6A**). The prevalence of these three main areas of effect was also independently highlighted by manual literature-based categorisation of gene functions.

**Figure 6:**
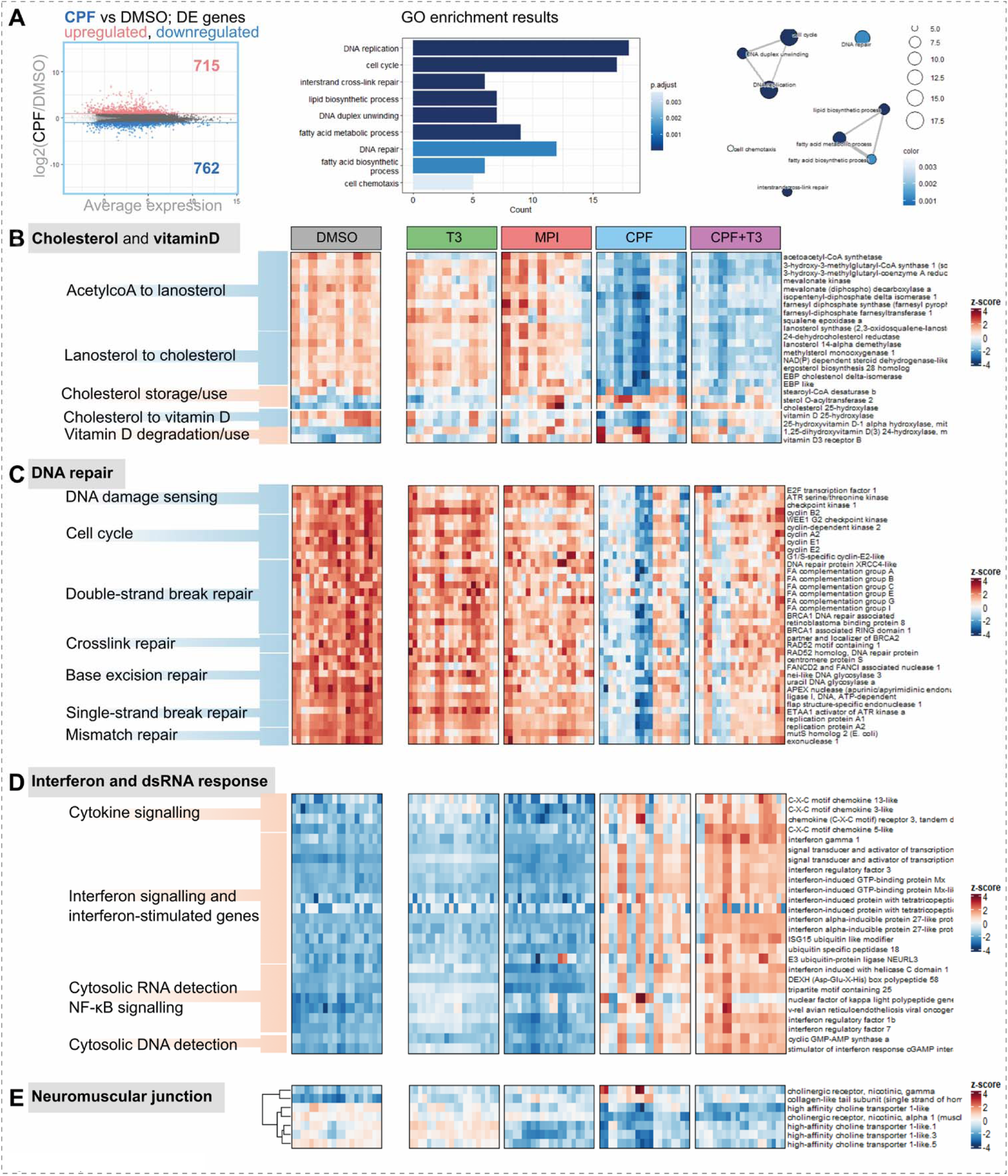
CPF affects DNA repair, immunity, as well as the cholesterol and vitaminD synthesis pathways. **A) Left:** MA plot of the log_2_ change in gene expression between CPF-treated larvae and vehicle-controls (DMSO), against the average expression level of each gene. Differentially expressed genes (abs(log_2_FC) ≥ 1; adj.p.val < 0.05) are coloured in red (upregulated) or blue (downregulated). **Right:** results from GO-enrichment analysis for CPF differentially expressed genes. **B)** expression pattern (z-scores) of DE genes involved in cholesterol and vitamin D metabolism. **C)** expression pattern of DE genes involved in DNA repair. **D)** expression pattern of DE genes involved interferon and dsRNA response. **E)** expression pattern of DE genes involved at the neuromuscular junction. For all heatmaps, each column within each treatment condition represents a different larva (with technical replicates). Gene rows in B) are ordered by metabolic pathway sequence; genes in C), D) are grouped by category; genes in E) are clustered based on pattern of expression.

Accordingly, we find that CPF-treated juveniles display a unique, coordinated, downregulation of nearly all genes involved in cholesterol synthesis (**Fig. 6B**). Specifically, and with few exceptions, all enzymatic functions leading from acetyl-CoA to the cholesterol final product are downregulated under CPF. Genes involved in cholesterol storage/export are upregulated (e.g. esterifying enzymes sterol O-acyltransferases) despite an overall downregulation of all genes encoding for apolipoproteins (see **Fig. S5C**). Our interpretation is that cells may be attempting to export as much as they can of the little cholesterol they have in the face of depressed cholesterol synthesis and cholesterol transport status. Notably, none of these metabolic changes can be rescued by T3 supplementation, and these pathways are not regulated by T3 or MPI alone. Moreover, the downregulation of the cholesterol synthesis pathway is accompanied by downregulation of the genes involved in vitamin D synthesis (liver vitamin D 25-hydroxylase *cyp2r1* and kidney 25-Hydroxyvitamin D 1-alpha-hydroxylase *cyp27b1*), matched by upregulation of vitamin D degradation (kidney vitamin D 24-hydroxylase *cyp24a1*). Notably, no other cholesterol-utilising pathway, notably steroidogenesis and bile acid conversion, undergoes analogous coordinated changes in expression (**Fig. S5**).

Another important set of targets downregulated by CPF, picked up by GO-driven analysis, involves DNA damage repair genes (**Fig. 6C**), consistent with the genotoxic effect of this pesticide observed in other contexts (e.g. Li et al., 2015). Of these, we specifically note the downregulation of multiple genes involved in the detection and repair of double-strand DNA breaks: most of the core Fanconi Anemia complex genes, the gene coding for the downstream FancI/KIAA1794 protein, both BRCA genes (Breast cancer 1 and 2 *brca1/2*) and their associated proteins (Retinoblastoma-binding protein 8 *rbbp8/CtIP*, BRCA1-associated RING domain protein 1 *bard1*, Partner and localizer of BRCA2 *palb2*), as well as the repair protein RAD52 homolog *rad52*. We also note that CPF exposure leads to downregulation of many base excision repair pathway components, including notably the Apurinic/apyrimidinic endodeoxyribonuclease 1 *apex1/ape1* (the major AP endonuclease of this pathway) and downstream repair enzymes DNA ligase I *lig1* and Flap structure-specific endonuclease 1 *fen1*, as well as the two DNA glycosylases Nei endonuclease VIII-like 3 *neil3* and Uracil-DNA glycosylase *unga*, detecting oxidised and deaminated bases respectively. Finally, we also detect downregulation of ssDNA damage detection proteins (Replication protein A 1 and 2, *rpa1/2*) as well as key ssDNA damage checkpoint components Serine/threonine-protein kinase ATR *atr* and Checkpoint kinase 1 *chek1*. Incidentally, CPF also leads to the downregulation of most of the major cyclins and cyclin-dependent kinases (cyclinB2; wee1, cdk2, cyclinA2/E1/E2; **Fig. 6C**), which are also key components of cell cycle checkpoint mechanisms. Again, none of these genes are differentially expressed by T3 or MPI alone. In summary, we detect transcriptomic responses suggesting that clownfish larvae exposed to CPF, or at least specific subpopulations of cells within them, are left extremely vulnerable to genomic instability (given the depressed repair capabilities) if not cell-cycle impaired. Downregulation of DNA repair genes in the absence of cell proliferation has been linked — in other contexts — to the activation of programmes of cellular senescence in response to genotoxic stress (Collins at al., 2018).

Among upregulated genes, a striking unique signature of CPF treatment consists in the upregulation of genes normally associated with antiviral response, or response to intracellular nucleic acids, and therefore also associated with a proinflammatory status (**Fig. 6D**). Apart from the upregulation of multiple cytokine and chemokine genes, pathway components of the type II interferon response (Interferon gamma 1 *ifng1*, Signal transducer and activator of transcription 1a and 2 *stat1a*, *stat2*), and a multitude of interferon response genes; we were surprised to see upregulation of pathways normally responsive to cytosolic viral RNAs and DNA. Specifically, we detect upregulation of both RIG (DEXH (Asp-Glu-X-His) box polypeptide 58 *dhx58*) and MAD5 signalling pathways (interferon induced with helicase C domain 1 *ifih1*) detecting ssRNA, short dsRNA, and long dsRNA; downstream transcriptional interferon regulatory factors, as well as multiple genes involved in the innate antiviral response or viral RNA translation termination. The dsRNA sensor zinc finger, NFX1-type containing 1 *znfx1* (Wang et al., 2019) is also strongly upregulated. Moreover, all components of the pathway detecting cytosolic dsDNA, cGAs-STING, are also uniquely upregulated in response to CPF treatment.

Interestingly, these effects seem to specifically affect Nk-κB components involved in the non-canonical signalling pathway (*relB + NfkB2*). These transcriptomic responses would normally be indicative of viral infection. We postulate that these pathways may be activated by the cell’s own nucleic acids, possibly in relation to the genomic instability detected above. In this case again, the transcriptomic response cannot be rescued by T3 supplementation and in fact appears to become more consistent across T3-treated juveniles.

Finally, because one of CPF’s main mechanism of action is the inhibition of neuromuscular junction cholinesterases (see e.g. Reiss et al., 2012), demonstrated at equivalent concentrations in juveniles of other reef species (Botté et al., 2012), we also tested whether we could recover such an effect in clownfish larvae, based on the pattern of expression of neuromuscular junction components (**Fig. 6E**). Accordingly, we observe transcriptomic changes consistent with cholinergic overstimulation and muscle damage, as also observed histologically in other CPF-treated fish (Sudhakaran et al., 2023). Clownfish larvae exposed to CPF downregulate a number of choline transporters (*slc5a7*-like) as well as the alpha subunit characteristic of the mature nicotinic acetylcholine receptor (*chrna1*). Similarly, the upregulation of the ColQ subunit of the acetylcholinesterase itself (*colq*) may suggest a compensatory attempt to increase recruitment of the enzyme at the neuromuscular junction due to its effective loss of activity under CPF. Interestingly, the neuromuscular damage experienced by larvae exposed to organophosphate is translated into the upregulation of the gamma subunit of the acetylcholine receptor (*chrng*), establishing channels with lower conductance typical of the receptors on foetal and denervated muscles (Mishina et al., 1986).

Overall, and in addition to the expected dysfunctionalisation of neuromuscular junction signalling, we find that CPF has major disruptive effects on the biology of metamorphosing larvae, mostly affecting the cholesterol synthesis pathway (and intermediate products), precursor of vitamin D. CPF also appears to induce a surprising immunological response, and drastic depression of the larval genomic repair machinery.

## 4. Discussion

Previous efforts in larval rearing development by our lab have succeeded in establishing low-volume rearing conditions for pre- and peri-metamorphosis clownfish stages (Roux et al., 2021). These modified rearing conditions allow us to perform routine pharmacological perturbation of clownfish larvae and early juveniles. By coupling such treatments with i) photographic documentation of the end-phenotype of each treated larva, and ii) end-of-treatment whole-larva RNA extraction + bulk RNA sequencing, we were able to systematically obtain peri-metamorphosis “treatment → phenotype → transcriptome” integrated datasets across a variety of pharmacological perturbations. We have adopted this approach to outline the response of *Amphiprion ocellaris* larvae to CPF during metamorphosis, and further use this transformation as a readout of TH activity to focus on the intersection between CPF and the TH-signalling pathway. We show that CPF exposure, by reducing TH levels, impairs the formation of white bands in clownfish larvae in a dose-dependent way, and alters the trajectory of larval growth, both readouts of metamorphosis progression. Interestingly these effects can be rescued by TH treatment, establishing a direct causal link between CPF effects and TH disruption. By transcriptomic analysis, we fully detail the effects of the pesticide (and T3 rescue) on the larval TH signalling pathway and find that CPF acts on this pathway as the reference goitrogen mix MPI. Still, we find that CPF also induces systemic, TH-independent, effects on cholesterol and vitamin D metabolism, DNA repair, and immunity, highlighting broader impacts on metamorphosing larvae that may not be immediately discernible from phenotypic analysis alone. Our findings thus explore two major areas of interest: 1) the extent- and the nature of the intersection between exposure to CPF, a potential endocrine disrupting compound, and the TH-synthesis axis, with potential applications on human and animal health; and 2) the general effects of CPF, as pesticide polluting many reef environments, on clownfish larvae undergoing a key transition period in their life, with implication on conservation, ecosystem management, and on studies of metamorphosis itself that in fact also extend to species that cannot be studied in a laboratory setting (e.g. surgeonfish; Holzer et al., 2017; Besson et al., 2020).

### 4.1. Endocrine disrupting effects of CPF on the TH-signalling axis

By profiling the response to CPF of all major- and accessory components of the TH-signalling pathway, we show that this pesticide acts *de facto* as the mix of goitrogens MPI on metamorphosing clownfish larvae, which we see translated in decreased measured T3 hormone levels. In addition to the observed downregulation of *duox1/2* and *dio1*, our results also highlight gene expression changes suggestive of hypothyroidism, as observed zebrafish larvae exposed to the organophosphate pesticide EHDPP (Shu et al., 2024). Though CPF has been described as being able to damage thyroid follicle cells (De Angelis et al., 2009), our CPF+T3 rescue experiments suggests that this is not the case in metamorphosing larvae (or is reversible), as normal function can be fully restored even in the presence of CPF. We were however unable to recover expression of some of the components of the HPT axis (*tsh*, *tshR*), likely also due to the minimal proportion of pituitary tissue in our samples. Consequently, we are unable to ascertain whether the suppressive action that we see on the TH signalling axis and on measured T3 levels may come from upstream disruption of hypothalamic or pituitary function (as suggested for CPF in zebrafish, and for other organophosphate pesticides in rat; Qiao et al., 2021; Xiong et al., 2018). Interestingly, Shu et al, 2024 show that the organophosphate pesticide EHDPP exerts its mechanism of action by competitively binding transthyretin (*ttr*) as a T4 mimic, such that the exact mode of action of CPF remains unclear. Our data suggests that the functional intersection between CPF and TH signalling lies upstream to T3 production, given our rescue experiment results and hormone measurement levels. Our data also suggests that the action of the pesticide does not derive from damage or irreversible inhibition of the cellular or molecular components of the TH synthesis and signalling axis, given that a T3-like TH-signalling signature can be re-established in the CPF-T3 condition even though CPF is still present.

Though we find that CPF has MPI-like effects at both the phenotypic, hormonal, and TH-signalling levels, we also note that only 36% of MPI regulated genes are also regulated by CPF. Given that MPI specifically targets the TH-signalling pathway, other compounds that have an MPI-like effect on this pathway would be expected to recover the full response signature of the goitrogen mix. The TH-signalling inhibitory effects of CPF instead do not entirely extend to all expected responsive genes. As summarised in **Fig. 7**, we expect any downstream effects of CPF’s goitrogenic action on the TH-signalling axis to have to occur within a widely dysfunctional, CPF-toxified context (i.e. a context of promiscuous acetylcholinesterase inhibition), which as such may only allow those goitrogen-induced effects possible in such a broadly altered context. Accordingly, our results may not be as unexpected: an underlying toxic effect of CPF on specific cell types or tissues of the clownfish larva may explain differences in the sets of responding TH-controlled genes despite the same TH-signalling context.

**Figure 7:**
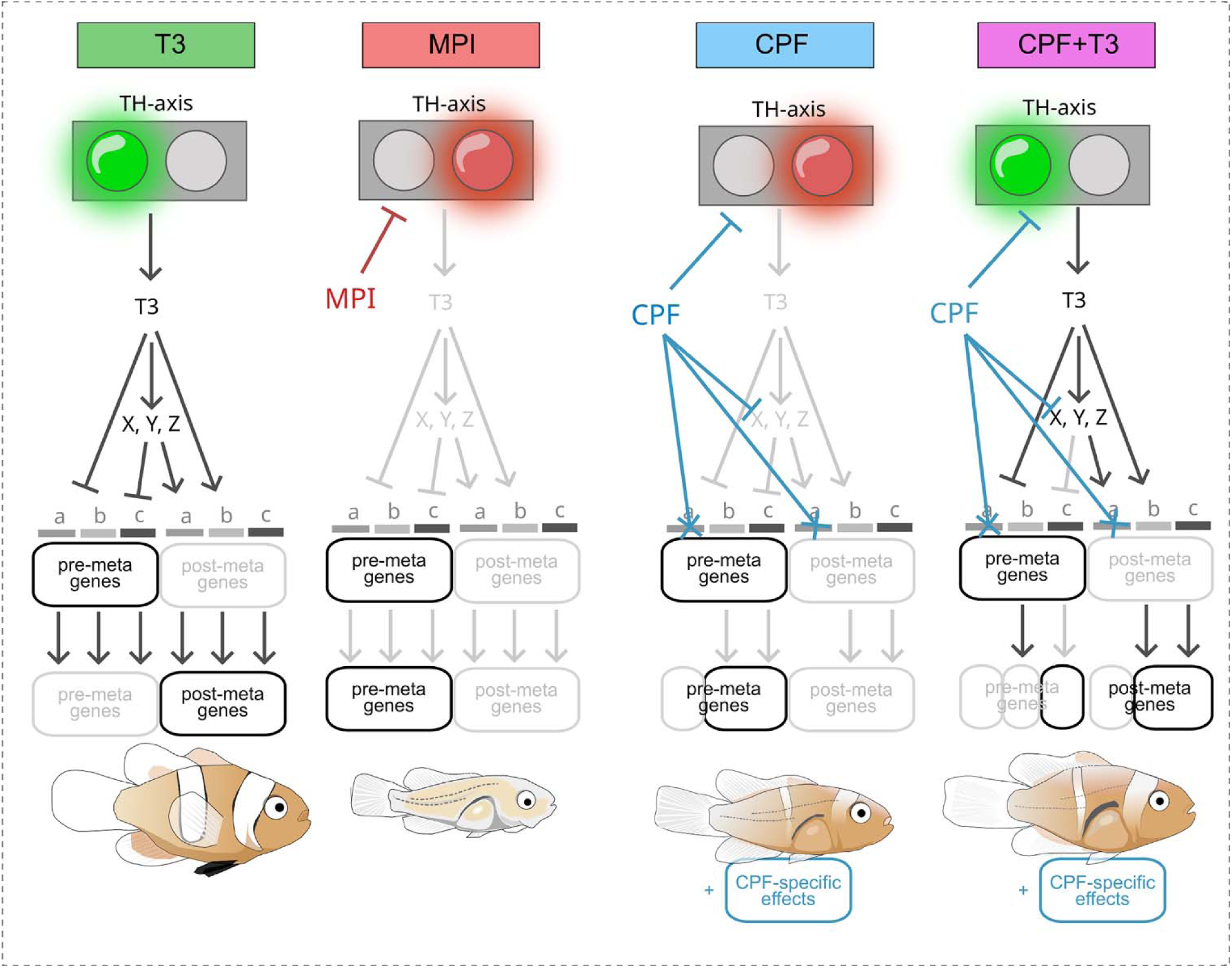
summary diagram of the intersection between CPF and TH-driven metamorphic changes, and the expected overlaps between post-metamorphosis gene expression profiles. The status of the TH axis (active or inactive, green or red light) determines T3 levels and its peripheral signalling effects on different larval tissues (“a”, “b”, “c”), through direct activation or repression, or in cooperation with intermediate co-effectors (“X”, “Y”, “Z”). During T3-coordinated metamorphosis, this leads to a switch between the expression of pre-metamorphosis genes (top row) to post-metamorphosis genes (bottom row), which would include, for example, the genes responsible for band pattern formation. The known and postulated actions of MPI and CPF, respectively, are illustrated. Notably, the incomplete overlap between MPI and CPF responses, despite both treatments leading to the same TH-signalling status, suggests that CPF affects either intermediary effectors, or prevents gene expression changes e.g. by toxic effects on specific tissues (here, tissue “a”). Note, secondary TH-control effectors repressed by T3, which normally block the expression of post-metamorphosis genes (and possibly activate the expression of pre-metamorphosis genes), are not shown for the sake of simplicity.

As discussed in the main body of the paper, our observation that CPF also shares 30% of the transcriptomics signatures of T3 treatment (i.e. TH–signalling stimulation) appears counterintuitive. Yet, by analysing these shared signatures we find clear examples illustrating that these may generally represent genes that, under T3, are downregulated as part of coordinated metabolic and physiological switches preparing the larva to its new post-metamorphosis environment, and, under CPF, are downregulated though clearly not as part of global compensatory changes in other gene sets. Taking the example of haemoglobin genes mentioned previously, a toxic effect of CPF on blood cells (e.g. tissue ’’a’’ in **Fig. 7**), would lead to observed downregulation of haemoglobin genes compared to control, and as such overlap with the coordinated downregulation of pre-metamorphosis haemoglobins in T3-treated larvae. In CPF however, post-metamorphosis haemoglobins are also downregulated, a signature that would instead be recovered in MPI-treated larvae. With the examples of the lipogenesis to beta-oxidation switch and of the haemoglobin complement switch, our results also incidentally underscore a key role of THs in the control of metamorphosis. That is, the importance of THs is not necessarily to be found in the specific genes downregulated or upregulated — as these same effects will also be recapitulated e.g. by pesticides — but in the fact that these effects are coordinated in time and at the system level to other changes in complementary sets of genes. Simply recapitulating either, without coordination, will not work.

### 4.2. Systemic effects of CPF on clownfish larvae

Beyond the TH-suppressive role of CPF, we further highlight three main domains of effect of the pesticide on clownfish larvae: downregulation of cholesterol and vitamin D synthesis genes, downregulation of of DNA damage/repair genes, and activation of immunity/antiviral genes.

The liver is the main site of cholesterol synthesis and CPF is known to have hepatotoxic effects in several organisms (Das et al., 2008; He et al., 2015; Akhtar et al., 2009) as it is a major site of bioactivation of the pesticide into its more dangerous derivative oxon (Sultatos 1991). Yet, the absence of change we detect in liver functions other than cholesterol synthesis, and the continued detected expression of other liver function genes, suggests that the downregulation of cholesterol synthesis genes we here describe is the result of a regulatory change rather than that of tissue cytotoxicity. Where investigated, CPF has been found in fact to *increase* cholesterol and triglyceride levels (He et al., 2015; Akhtar et al., 2009; Tanvir et al., 2016; Djekkoun e tal. 2022) through mechanisms opposite to the changes we detect here. Less is known for fish, though available data critically highlights the reverse effect: CPF often tends to result in lower measured cholesterol levels (Hatami et al., 2019; Ghayyur et al., 2019; Sanden et al; 2018), though contrasting reports exist (Abdel-Daim et al., 2020; Prakash 2020; Velmurugan et al., 2022). The response we see in clownfish larvae thus appears to represent a fish-specific response to CPF, with widespread implications given the central role played by cholesterol and its derivatives in animal biology. In terms of lipid metabolism, the selective downregulation of fatty acid desaturases under CPF matches quantitative analyses performed in CPF-exposed salmons, showing relative accumulation of saturated intermediates and a decrease in polyunsaturated fatty acids (Olsvik et al., 2015; Sanden et al; 2018). More literature at the CPF-cholesterol intersection focuses instead on cholesterol as precursor of corticosteroids, signalling components of another key endocrine pathway. Effects of CPF on the hypothalamic-pituitary-gonadal axis and on steroidogenic pathways indeed abound (Oruç 2010; see Ur Rahman 2021). In our study however, CPF did not seem to affect cholesterols-utilising pathways other than vitamin D metabolism. We note that the gonadal system in clownfish larvae is not developed, and in fact clownfish do not possess a differentiated gonad until much later in development (Miura et al., 2007), warning against extrapolating these data as a support to the absence of endocrine disrupting effects of CPF on steroidogenic and sexual differentiation endocrine systems. We also note that it is believed that vitamin D synthesis from endogenous cholesterol does not take place in fish (skin) due to this requiring a light-mediated step that may not be possible given the low penetrance of light through water (Lock et al., 2010). Yet, the coordinated shift we observe in both pathways (and not in other cholesterol branches), combined with experimental evidence of endogenous vitamin D synthesis in other fish species (Pierens et al., 2015), would suggest that clownfish larvae may in fact be capable of endogenous vitamin D synthesis, and that such a process is strongly impaired/downregulated following CPF exposure. Demonstration of effective vitamin D synthesis in clownfish larvae and quantification of vitamin D levels in response to CPF exposure will be needed to verify our inferences based on the transcriptomic response we detected.

The second major CPF effect we find signatures of — genotoxicity — has been described in several contexts and though contrasting results exist on the ability of the pesticide to actually bind DNA and create adducts (Cui et al., 2006; ur Rahman et al., 2021), it has been shown to induce both single-stranded and double-stranded DNA breaks in a variety of species and experimental contexts (Li et al, 2015; Yin et al., 2009; Ali et al., 2008, Mehta et al., 2008; Soum et al., 2022; see also ur Rahman et al., 2021), likely through the associated generation of reactive oxygen species (Gupta et al., 2010). In fact, Wang et al., 2024 see transgenerational effects of CPF exposure, which they link to the genotoxifying action of the pesticide. Critically, studies have suggested that CPF-exposure can cause not only damage of DNA, but also damage of its repair enzymes by reactive oxygen species and impair DNA metabolic functions (including repair) due to the induction of DNA-protein cross-links (Ohjia et al., 2015). We find a similar response, at the transcriptional level, in clownfish larvae: many of the larvae exposed to CPF undergo a strong downregulation of components involved at all levels of the DNA-repair apparatus, from DNA damage sensing to single strand and double strand break repair, to nucleotide excision repair. Clownfish larvae thus appear to become extremely sensitised to genomic instability (given the depressed repair capabilities) and these changes seem to be further coupled to the downregulation of cyclins and to the suppression of major cell cycle checkpoint effectors, suggesting cell cycle arrest. In fact, the signature we recover has been defined as a hallmark of cellular senescence (Collins et al., 2018). The question remains on whether this is a systemic response of the whole clownfish larva, or only of specific subpopulations within it. The fact that the genotoxic effects of CPF have often been observed in blood populations (e.g. Samajdar et al., 2015), and that we find CPF to be strongly toxifying to clownfish blood cells or to their precursors (downregulation of all haemoglobin genes), may suggest that it is blood populations that are the ones damaged the most. In fact, acetylcholinesterase is an important membrane component of red blood cells and erythropoietic cells (Lawson and Barr, 1987; Basirun et al., 2019) and, at least in humans, has been found to be a more sensitive readout of organophosphate exposure than other isoforms (Chen et al., 1999).

Finally, while the general immune stimulation triggered by CPF in clownfish larvae is not surprising given the oxidative damage expected from the pesticide and has indeed been observed across exposed fish species (e.g. Chen et al., 2014; Jin et al., 2015; reviewed in Díaz-Resendiz et al., 2015), the specific upregulation of innate immune response genes responsive to cytosolic nucleic acid, including dsDNA, is puzzling. To our knowledge, this response has not been described before and would be expected from viral infection. We note that this possibility is unlikely in our experimental setup, given that the antiviral response is specific to CPF-treated larvae, and all larvae in all treatments shared the same water source and the same air. It is however possible that CPF weakens clownfish larvae making them more susceptible to ambient viruses (natural sea water). Larvae in other treatment conditions would instead remain resistant to these viruses. Immunosuppressive roles of CPF are known (El-Bouhy et al., 2016), and the cytokine imbalance created by ongoing inflammatory processes (Chen et al., 2014) may itself be a source of sensitisation. A likely alternative explanation would be that the activation of viral-response-like pathways we observe is triggered not by viruses but by DNA leaking from damaged mitochondria. Oxidative damage (a key consequence of CPF biotransformation) has indeed been found to trigger pyroptosis, cause extracellular leakage of mitochondrial DNA and evoke cGAS-STING responses in other contexts (Miao et al., 2024).

### 4.3. Pesticides and larval metamorphosis (TH as a tuning dial)

We have recently elaborated an Eco-Evo-Devo model of the role of THs across the animal kingdom, whereby THs help animals one one hand to match their ontogenetic transitions with environmental conditions, and on the other hand to compensate adverse environmental conditions in order to nonetheless guarantee the successful completion of such transitions (see Zwahlen et al., 2024). In the context of this model, the results we describe in this paper highlight criticalities on the ecological consequences of organophosphate and pesticide pollution which we find of fundamental importance.

Specifically, the rescue experiments are particularly informative as they suggest that clownfish larvae could, for the most part, counter the effects of environmental pesticides (at least, of CPF) by a compensatory increase of T3 levels. Accordingly, supplementation of T3 to CPF leads to the recovery — at the scale of the whole transcriptome — of almost 70% of the T3-only signature, including those expression programmes that drive key transformations such as body shape changes and white band formation. This is a striking recovery effect, that highlights the considerable compensatory power held by THs as guardians of larval quality in the face of environmental challenge. Critically however, CPF also affects the very larval ability to synthesise T3, and thus nullifies any possibility for such compensatory changes to occur (that is, outside of a laboratory setting where such hormone could be supplied exogenously). We therefore put forward that pesticides with thyromodulatory potential, such as CPF, represent a qualitatively different kind of threat to marine larvae, and one of much bigger importance: aside from their systemic effects on larval physiology, these pesticides also critically block a major mechanism through which larvae could compensate such changes, as we reveal by supplying T3 exogenously.

Still, we note that were larvae able to produce more T3 and recover a T3-like gene expression signature, and thus were they able to metamorphose (CPF+T3, or the case of a non-thyromodulatory pesticide), they would be left with all the additional pesticide effects we have found to be TH-independent (for CPF: cholesterol repression, inflammation, genomic instability, neuromuscular toxicity). Pesticide pollution, regardless of its thyromodulatory effects, thus profoundly alters the fitness landscape of exposed larvae, such that visual readouts of successful metamorphosis may not be sufficient to assess the true extent of such alterations.

### 4.4. Technical, and limitations

CPF levels in coastal environments and reef ecosystems have been found to range from <3.6-83ng/L (Carvalho et al, 2002), to up to 0.5µg/L, sometimes exceeding 10µg/L, in Australian surface waters (NRA, 2000), with effects on fish larvae already at 0.5-1µg/L (Jarveinen et al., 1983; Besson et al., 2017; Botté et al., 2012; Besson et al., 2020; Bertucci et al., 2018). In this study we expose clownfish larvae to the higher end of such pesticide concentrations, with nominal concentrations ranging from 10µg/L to 30µg/L, measuring transcriptional responses at 20µg/L. These concentrations remain below those inducing behavioural abnormalities in metamorphic or juvenile stages of other reef fish (Besson et al., 2017; Botté et al., 2012) and fall within the range of what typically applied for CPF studies on these species (Besson et al., 2017; Botté et al., 2012; Holzer et al., 2017; Besson et al., 2020). We use relatively high and sublethal CPF concentrations to ensure we magnify effects that may be otherwise too weak to detect, and to be sure to capture the potential effects of the pesticide on the TH signalling axis, one of the primary aims of our investigation. Indeed, reports do not consider CPF as an endocrine disruptor at the concentrations that already cause acetylcholinesterase inhibition (Juberg et al., 2013), underscoring the fact that such effects may only become apparent at higher concentrations. Our lowest treatment concentration matches CPF’s IC_50_ (9.7 µg/L) for the inhibition of acetylcholinesterase activity in spiny damselfish juveniles (Botté et al., 2012), as well as the minimal concentrations required to see changes in TH levels in surgeonfish (Holzer et al., 2017; Besson et al., 2020). Though analogous studies at environmental concentrations will reveal whether these are already sufficient to disrupt larval development in wild clownfish larvae, the clear TH-disrupting, metamorphosis-impairing, and TH-independent systemic effects we here describe spell clear warnings about the environmental risk carried by this pollutant, underscoring the importance of its management. At the same time, it should also be noted that exposure to high-concentrations of CPF is a present reality for e.g. frontline agricultural workers using this pesticide (Liem et al., 2021), that TH-disrupting effects are noted across these demographics (Liem et al., 2023), and that concentrations of chlorpyrifos of 10-30 µg/L are already detected in waterways across the globe (Marino et al., 2005; Leong et al., 2007, Alvarez et al., 2009).

From a strictly Eco-Devo point of view, exposing clownfish larvae to high CPF levels also allows us to stress-test the buffering role we believe TH plays during post-embryonic larval development (as discussed above), and to assess the limits to which TH-mediated changes can effectively compensate changes in ecological conditions. Incidentally, even at these high CPF levels, we see that larval transcriptomic and hormonal responses are highly heterogeneous, hinting at the fact that such a heterogeneity may be a feature of the response to pesticide exposure, rather than a sign of underdosing. The observed heterogeneity in response is similarly reflected in our TH-level measurements, though this may be exacerbated by the limited number of replicates we had available for hormone assays and confounded by the need to use pools of larvae, obscuring individual responses to CPF exposure. Still, the larval heterogeneity we observe (genetic or other) may itself be an adaptive strategy of marine species, favouring resilience to pollutant challenge just as it is starting to be recognised in the context of fish climate change resilience strategies (Schunter et al., 2022). Overall, our results enhance understanding of the intricate interplay between CPF exposure, TH signalling and metamorphosis, emphasising the urgent need for mitigating the detrimental consequences of chemical pollutants on marine ecosystems.

## Data Availability

- Raw reads of bulkRNAseq datasets are available to download under NCBI BioProject accession number PRJNA1138483 (https://www.ncbi.nlm.nih.gov/bioproject/PRJNA1138483)
- Counts matrix and code to reproduce the transcriptomics analysis is available at: https://github.com/StefanoVianello/ReynaudVianello_AoceCPF. Lists of all genes in each treatment intersection, as indicated in the text, are also made available.

### Acknowledgments

We thank the High Throughput Genomics Core of the Biodiversity Research Center at Academia Sinica for performing the NGS experiments. The core facility is funded by Academia Sinica Core Facility and Innovative Instrument Project (AS-CFII-108-114). We further thank Dr. Mei-Yeh Lu and Ms. Pei-Lin Chao for their assistance and discussion troubleshooting quality-control metrics. We also thank the staff of the ICOB Marine Research Station for superb help for fish husbandry. Work from our laboratory is supported by a Grand Challenge Grant from Academia Sinica and JSPS KAKENHI grant 22H02678 at OIST. L.B. and D.L. have been supported by a grant from Agence Nationale de la Recherche SENSO (ANR19-CE14-0010).

## Supporting information: figure captions

***Figure S1: The white band phenotype at 19 dpt***

*Percentage of larvae having 0 (white), 1 (yellow), 2 (orange) or 3 (dark orange) white bands. (nDMSO=21, nCPF 20µg/l =11, n MPI= 16, nT3 = 21 individuals).*

***Figure S2: anatomical landmarks of geometric morphology approach.***

*Landmarks (in red) used in this study defining the overall body shape of Amphiprion ocellaris shown at two stages: 0 dpt (left) and 5 dpt (right). The landmarks are as follow: (1) centre of the eye; (2) mouth tip; (3) end of lower jaw articulation; (4) anterior insertion of the stomach; (5) anus; (6) posterior insertion of anal fin; (7) ventral base of the caudal fin; (8) end of notochords (9) posterior insertion of the anal fin; (10) anterior insertion of the anal fin; (11) dorsal insertion of the stomach, (12) posterior end of the nuchal crest; (13) dorsal margin through the midline of the eye. Ten curve landmarks (in orange) have been used to specifically capture the head shape changes.*

***Figure S3: Phenotype of treated larvae used for bulkRNA seq.***

*Photos of larvae after 5 days of treatment under each of the conditions indicated (final age: 13dph). From top to bottom: DMSO vehicle control, T3, MPI, CPF, and CPF+T3. For 10 larvae from each treatment total RNA was extracted following picture acquisition (except larvae with red strike through; see materials and methods). Larvae framed in yellow: CPF-treated larvae with strongest downregulation of iridophore genes compared to control; see text relating to Fig. 5). Scale bar: 1mm.*

***Figure S4: Comparative overview of the pattern of expression of differentially expressed genes under each treatment.***

*For each of the three single treatments T3 (**A**), MPI (**B**), CPF (**C**): **Left**: “MA” plots (log_2_ change in gene expression between treated larvae and vehicle-controls (DMSO), against the average expression level of each gene). Differentially expressed genes are highlighted in blue and red (abs(log_2_FC))>1; adj.p.Val < 0.05). **Right**: expression pattern (z-scores) of upregulated (top) and downregulated (bot-tom) differentially expressed genes under each treatment. Asterisk: single treatment with the closest expression pattern (see materials and methods). Green: T3; Red: MPI; Blue: CPF. Responses unlike either of the two other treatments are in grey. Responses similar to both other treatments are in white (no colour). The MA plot in (C) is shown again in Fig. 6.*

***Figure S5: CPF effects on other gene sets of interest.***

*Expression pattern (z-scores) of genes involved in other liver- and cholesterol-related pathways. All genes shown have different expression in CPF than control, at adj.p.value < 0.05. Genes highlighted in blue have abs(logFC)>1 (i.e. they pass threshold of differential expression). For all heatmaps, each column within each treatment condition represents a different larva (with technical replicates).*

## Bibliography

Abdel-Daim, Mohamed M., Mahmoud AO Dawood, Mohamed Elbadawy, Lotfi Aleya, and Saad Alkahtani. "Spirulina platensis reduced oxidative damage induced by chlorpyrifos toxicity in Nile tilapia (Oreochromis niloticus)." Animals 10, no. 3 (2020): 473.

Abreu-Villaça, Y., & Levin, E. D. (2017). Developmental neurotoxicity of succeeding generations of insecticides. Environment International, 99, 55–77. 10.1016/J.ENVINT.2016.11.019

Adams, D. C., & Otárola-Castillo, E. (2013). Geomorph: An r package for the collection and analysis of geometric morphometric shape data. Methods in Ecology and Evolution, 4(4), 393–399. 10.1111/2041-210X.12035

Akhtar, Nahid, M. K. Srivastava, and R. B. Raizada. "Assessment of chlorpyrifos toxicity on certain organs in rat, Rattus norvegicus." J Environ Biol 30, no. 6 (2009): 1047–1053.

Ali, Daoud, Naresh Sahebrao Nagpure, Sudhir Kumar, Ravindra Kumar, and Basdeo Kushwaha. "Genotoxicity assessment of acute exposure of chlorpyrifos to freshwater fish Channa punctatus (Bloch) using micronucleus assay and alkaline single-cell gel electrophoresis." Chemosphere 71, no. 10 (2008): 1823–1831.

Alvarez, Melina, Cecile Du Mortier, Soledad Jaureguiberry, and Andrés Venturino. "Joint probabilistic analysis of risk for aquatic species and exceedence frequency for the agricultural use of chlorpyrifos in the Pampean region, Argentina." Environmental toxicology and chemistry 38, no. 8 (2019): 1748–1755.

Aman, A. J., Kim, M., Saunders, L. M., & Parichy, D. M. (2021). Thyroid hormone regulates abrupt skin morphogenesis during zebrafish postembryonic development. Developmental Biology, 477, 205–218. 10.1016/j.ydbio.2021.05.019

Andrews, S. (2010). FastQC: A Quality Control Tool for High Throughput Sequence Data [Online]

Baken, E. K., Collyer, M. L., Kaliontzopoulou, A., & Adams, D. C. (2021). geomorph v4.0 and gmShiny: Enhanced analytics and a new graphical interface for a comprehensive morphometric experience. Methods in Ecology and Evolution, 12(12), 2355–2363. 10.1111/2041-210X.13723

Bao, B., Ke, Z., Xing, J., Peatman, E., Liu, Z., Xie, C., Xu, B., Gai, J., Gong, X., Yang, G., Jiang, Y., Tang, W., & Ren, D. (2011). Proliferating cells in suborbital tissue drive eye migration in flatfish. Developmental Biology, 351(1), 200–207. 10.1016/j.ydbio.2010.12.032

Bartley, R., Waters, D., Turner, R., Kroon, F., Garzon-Garcia, A., Kuhnert, P., Lewis, S., Smith, R., Bainbridge, Z., Olley, J., Brooks, A., Burton, J., Brodie, J., & Waterhouse, J. (2017). 2017 Scientific Consensus Statement: land use impacts on the Great Barrier Reef water quality and ecosystem condition, Chapter 2: sources of sediment, nutrients, pesticides and other pollutants to the Great Barrier Reef. http://www.reefplan.qld.gov.au/about/assets/2017-scientific-consensus-statement-summary-chap02.pdf

Basirun, Ain Aqilah, Siti Aqlima Ahmad, Mohd Khalizan Sabullah, Nur Adeela Yasid, Hassan Mohd Daud, Ariff Khalid, and Mohd Yunus Shukor. "In vivo and in vitro effects on cholinesterase of blood of Oreochromis mossambicus by copper." 3 Biotech 9 (2019): 1–12.

Baumann, L., Ros, A., Rehberger, K., Neuhauss, S. C. F., & Segner, H. (2016). Thyroid disruption in zebrafish (Danio rerio) larvae: Different molecular response patterns lead to impaired eye development and visual functions. Aquatic Toxicology, 172, 44–55. 10.1016/j.aquatox.2015.12.015

Besson, Marc, William E. Feeney, Isadora Moniz, Loïc François, Rohan M. Brooker, Guillaume Holzer, Marc Metian, Natacha Roux, Vincent Laudet, and David Lecchini. "Anthropogenic stressors impact fish sensory development and survival via thyroid disruption." Nature Communications 11, no. 1 (2020): 3614.

Besson, Marc, Camille Gache, Frédéric Bertucci, Rohan M. Brooker, Natacha Roux, Hugo Jacob, Cécile Berthe, Valeria Anna Sovrano, Danielle L. Dixson, and David Lecchini. "Exposure to agricultural pesticide impairs visual lateralization in a larval coral reef fish." Scientific Reports 7, no. 1 (2017): 9165.

Bishop, C. D., D. F. Erezyilmaz, Thomas Flatt, C. D. Georgiou, M. G. Hadfield, A. Heyland, J. Hodin et al. "What is metamorphosis?." Integrative and Comparative Biology 46, no. 6 (2006): 655–661.

Blanton, Michael L., and Jennifer L. Specker. "The hypothalamic-pituitary-thyroid (HPT) axis in fish and its role in fish development and reproduction." Critical reviews in toxicology 37, no. 1-2 (2007): 97–115.

Bocquené, G. and Franco, A., 2005. Pesticide contamination of the coastline of Martinique. Marine Pollution Bulletin, 51(5-7), pp.612–619.

Bosu, Subrajit, Natarajan Rajamohan, Shatha Al Salti, Manivasagan Rajasimman, and Papiya Das. "Biodegradation of chlorpyrifos pollution from contaminated environment-A review on operating variables and mechanism." Environmental Research (2024): 118212.

Botté, E. S., D. R. Jerry, S. Codi King, C. Smith-Keune, and A. P. Negri. "Effects of chlorpyrifos on cholinesterase activity and stress markers in the tropical reef fish Acanthochromis polyacanthus." Marine pollution bulletin 65, no. 4-9 (2012): 384–393.

Brunson, Jason Cory. "Ggalluvial: layered grammar for alluvial plots." Journal of Open Source Software 5, no. 49 (2020).

Campinho, M. A. (2019). Teleost metamorphosis: The role of thyroid hormone. Frontiers in Endocrinology, 10(JUN), 383. 10.3389/fendo.2019.00383

Carvalho, F. P., Gonzalez-Farias, F., Villeneuve, J. P., Cattini, C., Hernandez-Garza, M., Mee, L. D., & Fowler, S. W. (2002). Distribution, fate and effects of pesticide residues in tropical coastal lagoons of northwestern mexico. Environmental Technology (United Kingdom), 23(11), 1257–1270. 10.1080/09593332308618321

Chen, Dechun, Ziwei Zhang, Haidong Yao, Ye Cao, Houjuan Xing, and Shiwen Xu. "Pro-and anti-inflammatory cytokine expression in immune organs of the common carp exposed to atrazine and chlorpyrifos." Pesticide biochemistry and physiology 114 (2014): 8–15.

Chen, William L., Joel J. Sheets, Richard J. Nolan, and Joel L. Mattsson. "Human red blood cell acetylcholinesterase inhibition as the appropriate and conservative surrogate endpoint for establishing chlorpyrifos reference dose." Regulatory toxicology and pharmacology 29, no. 1 (1999): 15–22.

Chen, Yunshun, Aaron TL Lun, and Gordon K. Smyth. "From reads to genes to pathways: differential expression analysis of RNA-Seq experiments using Rsubread and the edgeR quasi-likelihood pipeline." F1000Research 5 (2016).

Chen, Yunshun, Lizhong Chen, Aaron TL Lun, Pedro L. Baldoni, and Gordon K. Smyth. "edgeR 4.0: powerful differential analysis of sequencing data with expanded functionality and improved support for small counts and larger datasets." bioRxiv (2024): 2024–01.

Chopra, Kunal, Shoko Ishibashi, and Enrique Amaya. "Zebrafish duox mutations provide a model for human congenital hypothyroidism." Biology Open 8, no. 2 (2019): bio037655.

Collin, Guillaume, Anda Huna, Marine Warnier, Jean-Michel Flaman, and David Bernard. "Transcriptional repression of DNA repair genes is a hallmark and a cause of cellular senescence." Cell death & disease 9, no. 3 (2018): 259.

Collyer, M.L. and Adams, D.C., 2018. RRPP: An r package for fitting linear models to high dimensional data using residual randomization. Methods in Ecology and Evolution, 9(7), pp.1772–1779.

Cui, Yong, Jiangfeng Guo, Bujin Xu, and Ziyuan Chen. "Potential of chlorpyrifos and cypermethrin forming DNA adducts." Mutation Research/Genetic Toxicology and Environmental Mutagenesis 604, no. 1-2 (2006): 36–41.

Danecek, Petr, James K. Bonfield, Jennifer Liddle, John Marshall, Valeriu Ohan, Martin O. Pollard, Andrew Whitwham et al. "Twelve years of SAMtools and BCFtools." Gigascience 10, no. 2 (2021): giab008.

Das, Parikshit C., Yan Cao, Randy L. Rose, Nathan Cherrington, and Ernest Hodgson. "Enzyme induction and cytotoxicity in human hepatocytes by chlorpyrifos and N, N-diethyl-m-toluamide (DEET)." Drug metabolism and drug interactions 23, no. 3-4 (2008): 237–260.

De Angelis, Simona, Roberta Tassinari, Francesca Maranghi, Agostino Eusepi, Antonio Di Virgilio, Flavia Chiarotti, Laura Ricceri et al. "Developmental exposure to chlorpyrifos induces alterations in thyroid and thyroid hormone levels without other toxicity signs in Cd1 mice." Toxicological Sciences 108, no. 2 (2009): 311–319.

De Groef, Bert, Serge Van der Geyten, Veerle M. Darras, and Eduard R. Kühn. "Role of corticotropin-releasing hormone as a thyrotropin-releasing factor in non-mammalian vertebrates." General and comparative endocrinology 146, no. 1 (2006): 62–68.

de Jesus, Evelyn Grace, Yasuo Inui, and Tetsuya Hirano. "Cortisol enhances the stimulating action of thyroid hormones on dorsal fin-ray resorption of flounder larvae in vitro." General and Comparative Endocrinology 79, no. 2 (1990): 167–173.

Djekkoun, Narimane, Flore Depeint, Marion Guibourdenche, Hiba El Khayat El Sabbouri, Aurélie Corona, Larbi Rhazi, Jerome Gay-Queheillard et al. "Chronic perigestational exposure to chlorpyrifos induces perturbations in gut bacteria and glucose and lipid markers in female rats and their offspring." Toxics 10, no. 3 (2022): 138.

Downie, Adam T., Sjannie Lefevre, Björn Illing, Jessica Harris, Michael D. Jarrold, Mark I. McCormick, Göran E. Nilsson, and Jodie L. Rummer. "Rapid physiological and transcriptomic changes associated with oxygen delivery in larval anemonefish suggest a role in adaptation to life on hypoxic coral reefs." PLoS biology 21, no. 5 (2023): e3002102.

Durinck, Steffen, Paul T. Spellman, Ewan Birney, and Wolfgang Huber. "Mapping identifiers for the integration of genomic datasets with the R/Bioconductor package biomaRt." Nature protocols 4, no. 8 (2009): 1184–1191.

Durinck, Steffen, Yves Moreau, Arek Kasprzyk, Sean Davis, Bart De Moor, Alvis Brazma, and Wolfgang Huber. "BioMart and Bioconductor: a powerful link between biological databases and microarray data analysis." Bioinformatics 21, no. 16 (2005): 3439–3440.

Díaz-Resendiz, Karina Janice Guadalupe, Gladys Alejandra Toledo-Ibarra, and Manuel Ivan Girón-Pérez. "Modulation of immune response by organophosphorus pesticides: fishes as a potential model in immunotoxicology." Journal of immunology research 2015 (2015).

El-Bouhy, Z., G. El-Nobi, R. Reda, and R. Ibrahim. "Effect of insecticide ‘‘chlorpyrifos” on immune response of Oreochromis niloticus. Zagazig Vet. J. 44, 196–204." (2016).

Farías-Serratos, B. M., Lazcano, I., Villalobos, P., Darras, V. M., & Orozco, A. (2021). Thyroid hormone deficiency during zebrafish development impairs central nervous system myelination. PLOS ONE, 16(8), e0256207. 10.1371/journal.pone.0256207

Fortenberry, G. Z., Hu, H., Turyk, M., Barr, D. B., & Meeker, J. D. (2012). Association between urinary 3, 5, 6-trichloro-2-pyridinol, a metabolite of chlorpyrifos and chlorpyrifos-methyl, and serum T4 and TSH in NHANES 1999-2002. Science of the Total Environment, 424, 351–355. 10.1016/j.scitotenv.2012.02.039

Galindo, D., Sweet, E., DeLeon, Z., Wagner, M., DeLeon, A., Carter, C., McMenamin, S. K., & Cooper, W. J. (2019). Thyroid hormone modulation during zebrafish development recapitulates evolved diversity in danionin jaw protrusion mechanics. Evolution and Development, 21(5), 231–246. 10.1111/ede.12299

Gao, Chun-Hui, Guangchuang Yu, and Peng Cai. "ggVennDiagram: an intuitive, easy-to-use, and highly customizable R package to generate Venn diagram." Frontiers in Genetics 12 (2021): 706907.

Ghayyur, Shehzad, Sadia Tabassum, Munawar Saleem Ahmad, Naveed Akhtar, and Muhammad Fiaz Khan. "Effect of chlorpyrifos on hematological and seral biochemical components of fish Oreochromis mossambicus." Pakistan journal of zoology 51, no. 3 (2019): 1047.

Gu, Z., Eils, R., & Schlesner, M. (2016). Complex heatmaps reveal patterns and correlations in multidimensional genomic data. Bioinformatics, 32(18), 2847–2849.

Gu, Zuguang. Complex heatmap visualization. Imeta, 2022, vol. 1, no 3, p. E43.

Guillot, R., Muriach, B., Rocha, A., Rotllant, J., Kelsh, R. N., & Cerdá-Reverter, J. M. (2016). Thyroid Hormones Regulate Zebrafish Melanogenesis in a Gender-Specific Manner. PLOS ONE, 11(11), e0166152. 10.1371/JOURNAL.PONE.0166152

Gupta, Subash C., et al. "Chlorpyrifos induces apoptosis and DNA damage in Drosophila through generation of reactive oxygen species." Ecotoxicology and Environmental Safety 73.6 (2010): 1415-1423.

Hatami, Mahdiye, Mahdi Banaee, and Behzad Nematdoost Haghi. "Sub-lethal toxicity of chlorpyrifos alone and in combination with polyethylene glycol to common carp (Cyprinus carpio)." Chemosphere 219 (2019): 981–988.

Haynes, D., & Johnson, J. E. (2000). Organochlorine, Heavy Metal and Polyaromatic Hydrocarbon Pollutant Concentrations in the Great Barrier Reef (Australia) Environment: a Review. Marine Pollution Bulletin, 41(7–12), 267–278. 10.1016/S0025-326X(00)00134-X

He, Wei, Wenli Guo, Yi Qian, Shuping Zhang, Difeng Ren, and Sijin Liu. "Synergistic hepatotoxicity by cadmium and chlorpyrifos: disordered hepatic lipid homeostasis." Molecular medicine reports 12, no. 1 (2015): 303–308.

Holzer, G., Besson, M., Lambert, A., François, L., Barth, P., Gillet, B., Hughes, S., Piganeau, G., Leulier, F., Viriot, L., Lecchini, D., & Laudet, V. (2017). Fish larval recruitment to reefs is a thyroid hormone-mediated metamorphosis sensitive to the pesticide chlorpyrifos. ELife, 6. 10.7554/eLife.27595

Huerlimann, Roger, Natacha Roux, Ken Maeda, Polina Pilieva, Saori Miura, Hsiao Chian, Michael Izumiyama, Vincent Laudet, and Timothy Ravasi. "The transcriptional landscape underlying larval development and metamorphosis in the Malabar grouper (Epinephelus malabaricus)." eLife 13 (2024).

Islam, M. S., & Tanaka, M. (2004). Impacts of pollution on coastal and marine ecosystems including coastal and marine fisheries and approach for management: A review and synthesis. In Marine Pollution Bulletin (Vol. 48, Issues 7–8, pp. 624–649). Pergamon. 10.1016/j.marpolbul.2003.12.004

Jarvinen, Alfred W., and Danny K. Tanner. "Toxicity of selected controlled release and corresponding unformulated technical grade pesticides to the fathead minnow Pimephales promelas." Environmental Pollution Series A, Ecological and Biological 27, no. 3 (1982): 179–195.

Jason Cory Brunson and Quentin D. Read (2023). ggalluvial: Alluvial Plots in ’ggplot2’. R package version 0.12.5. http://corybrunson.github.io/ggalluvial/

Jin, Yuanxiang, Zhenzhen Liu, Tao Peng, and Zhengwei Fu. "The toxicity of chlorpyrifos on the early life stage of zebrafish: a survey on the endpoints at development, locomotor behavior, oxidative stress and immunotoxicity." Fish & shellfish immunology 43, no. 2 (2015): 405–414.

Juberg, D. R., Gehen, S. C., Coady, K. K., LeBaron, M. J., Kramer, V. J., Lu, H., & Marty, M. S. (2013). Chlorpyrifos: Weight of evidence evaluation of potential interaction with the estrogen, androgen, or thyroid pathways. Regulatory Toxicology and Pharmacology, 66(3), 249–263. 10.1016/J.YRTPH.2013.03.003

Keer, S., Storch, J. D., Nguyen, S., Prado, M., Singh, R., Hernandez, L. P., & McMenamin, S. K. (2022). Thyroid hormone shapes craniofacial bones during postembryonic zebrafish development. Evolution and Development, 24(1–2), 61–76. 10.1111/ede.12399

King, Juliette, Frances Alexander, and Jon Brodie. "Regulation of pesticides in Australia: The Great Barrier Reef as a case study for evaluating effectiveness." Agriculture, ecosystems & environment 180 (2013): 54–67.

Kitada, Y., Kawahata, H., Suzuki, A. and Oomori, T., 2008. Distribution of pesticides and bisphenol A in sediments collected from rivers adjacent to coral reefs. Chemosphere, 71(11), pp.2082–2090.

Kumar, S., Kaushik, G., & Villarreal-Chiu, J. F. (2016). Scenario of organophosphate pollution and toxicity in India: A review. Environmental Science and Pollution Research, 23(10), 9480–9491. 10.1007/s11356-016-6294-0

Langmead, Ben, and Steven L. Salzberg. "Fast gapped-read alignment with Bowtie 2." Nature methods 9, no. 4 (2012): 357–359.

Larsen, Donald A., Penny Swanson, Jon T. Dickey, Jean Rivier, and Walton W. Dickhoff. "In vitrothyrotropin-releasing activity of corticotropin-releasing hormone-family peptides in Coho Salmon, Oncorhynchus kisutch." General and Comparative Endocrinology 109, no. 2 (1998): 276–285.

Laudet, V. (2023). The multi-level regulation of clownfish metamorphosis by thyroid hormones. Cell Reports, 42(7). 10.1016/j.celrep.2023.112661

Laudet, Vincent, and Timothy Ravasi, eds. Evolution, development and ecology of anemonefishes: model organisms for marine science. CRC Press, 2022.

Laudet, V. (2011). The Origins and Evolution of Vertebrate Metamorphosis. Current Biology, 21(18), R726–R737. 10.1016/J.CUB.2011.07.030

Lawson, A.A. and Barr, R.D., 1987. Acetylcholinesterase in red blood cells. American journal of hematology, 26(1), pp.101–112.

Leis, J. M., Siebeck, U., & Dixson, D. L. (2011). How Nemo Finds Home: The Neuroecology of Dispersal and of Population Connectivity in Larvae of Marine Fishes. Integrative and Comparative Biology, 51(5), 826–843. 10.1093/icb/icr004

Leis, J. M., & Fisher, R. (2006). Swimming speed of settlement-stage reef-fish larvae measured in the laboratory and in the field: a comparison of critical speed and in situ speed Thesis-Spatial and temporal water quality changes during a large scale dredging operation View project Signif. https://www.researchgate.net/publication/280976006

Leong, Kok Hoong, LL Benjamin Tan, and Ali Mohd Mustafa. "Contamination levels of selected organochlorine and organophosphate pesticides in the Selangor River, Malaysia between 2002 and 2003." Chemosphere 66, no. 6 (2007): 1153–1159.

Li, Diqiu, Qingchun Huang, Miaoqing Lu, Lei Zhang, Zhichuan Yang, Mimi Zong, and Liming Tao. "The organophosphate insecticide chlorpyrifos confers its genotoxic effects by inducing DNA damage and cell apoptosis." Chemosphere 135 (2015): 387–393.

Libin, He, Huang Zhen, Wu Shuiqing, and Zheng Leyun. "Transcriptome analysis identifies candidate genes related to albinism mechanism in the skin of the Picasso clownfish." Acta Oceanographica 44, no. 2 (2022): 67–76.

Liem, Jen Fuk, Imam Subekti, Muchtaruddin Mansyur, Dewi S. Soemarko, Aria Kekalih, Franciscus D. Suyatna, Dwi A. Suryandari, Safarina G. Malik, and Bertha Pangaribuan. "The determinants of thyroid function among vegetable farmers with primary exposure to chlorpyrifos: A cross-sectional study in Central Java, Indonesia." Heliyon 9, no. 6 (2023).

Liem, Jen Fuk, Muchtaruddin Mansyur, Dewi S. Soemarko, Aria Kekalih, Imam Subekti, Franciscus D. Suyatna, Dwi A. Suryandari, Safarina G. Malik, and Bertha Pangaribuan. "Cumulative exposure characteristics of vegetable farmers exposed to Chlorpyrifos in Central Java–Indonesia; a cross-sectional study." BMC Public Health 21, no. 1 (2021): 1066.

Lock, E.J., Waagbø, R., Wendelaar Bonga, S. and Flik, G., 2010. The significance of vitamin D for fish: a review. Aquaculture nutrition, 16(1), pp.100–116.

Love, Michael I., Wolfgang Huber, and Simon Anders. "Moderated estimation of fold change and dispersion for RNA-seq data with DESeq2." Genome biology 15 (2014): 1–21.

Marino, Damián José Gabriel, and Alicia Estela Ronco. "Cypermethrin and chlorpyrifos concentration levels in surface water bodies of the Pampa Ondulada, Argentina." Bulletin of environmental contamination and toxicology 75 (2005).

McCarthy, Davis J., Yunshun Chen, and Gordon K. Smyth. "Differential expression analysis of multifactor RNA-Seq experiments with respect to biological variation." Nucleic acids research 40, no. 10 (2012): 4288–4297.

McMenamin, S. K., & Parichy, D. M. (2013). Metamorphosis in Teleosts. In Current Topics in Developmental Biology (Vol. 103, pp. 127–165). Academic Press. 10.1016/B978-0-12-385979-2.00005-8

Mehta, Anugya, Radhey S. Verma, and Nalini Srivastava. "Chlorpyrifos induced DNA damage in rat liver and brain." Environmental and molecular mutagenesis 49, no. 6 (2008): 426–433.

Miao, Zhiruo, Wei Wang, Zhiying Miao, Qiyuan Cao, and Shiwen Xu. "Role of Selenoprotein W in participating in the progression of non-alcoholic fatty liver disease." Redox Biology (2024): 103114.

Mishina, Masayoshi, Toshiyuki Takai, Keiji Imoto, Masaharu Noda, Tomoyuki Takahashi, Shosaku Numa, Christoph Methfessel, and Bert Sakmann. "Molecular distinction between fetal and adult forms of muscle acetylcholine receptor." Nature 321, no. 6068 (1986): 406–411.

Miura, Saori, Shigeo Nakamura, Yasuhisa Kobayashi, Francesc Piferrer, and Masaru Nakamura. "Differentiation of ambisexual gonads and immunohistochemical localization of P450 cholesterol side-chain cleavage enzyme during gonadal sex differentiation in the protandrous anemonefish, Amphiprion clarkii." Comparative Biochemistry and Physiology Part B: Biochemistry and Molecular Biology 149, no. 1 (2008): 29–37.

NRA, 2000. NRA Review of Chlorpyrifos, vol. 1, Series 00.5. National Registration Authority for Agricultural and Veterinary Chemicals.

Ojha, A., and Y. K. Gupta. "Evaluation of genotoxic potential of commonly used organophosphate pesticides in peripheral blood lymphocytes of rats." Human & Experimental Toxicology 34, no. 4 (2015): 390–400.

Olsvik, Pål A., Marc HG Berntssen, and Liv Søfteland. "Modifying effects of vitamin E on chlorpyrifos toxicity in Atlantic salmon." PLoS one 10, no. 3 (2015): e0119250.

Oruç, E.Ö., 2010. Oxidative stress, steroid hormone concentrations and acetylcholinesterase activity in Oreochromis niloticus exposed to chlorpyrifos. Pesticide biochemistry and physiology, 96(3), pp.160–166.

Otênio, J. K., Souza, K. D., Alberton, O., Alberton, L. R., Moreno, K. G. T., Gasparotto Junior, A., Palozi, R. A. C., Lourenço, E. L. B., & Jacomassi, E. (2022). Thyroid-disrupting effects of chlorpyrifos in female Wistar rats. Drug and Chemical Toxicology, 45(1), 387–392. 10.1080/01480545.2019.1701487

Patro, Rob, Geet Duggal, Michael I. Love, Rafael A. Irizarry, and Carl Kingsford. "Salmon provides fast and bias-aware quantification of transcript expression." Nature methods 14, no. 4 (2017): 417–419.

Pierens, S. L., and D. R. Fraser. "The origin and metabolism of vitamin D in rainbow trout." The Journal of steroid biochemistry and molecular biology 145 (2015): 58–64.

Polidoro, B. A., Comeros-Raynal, M. T., Cahill, T., & Clement, C. (2017). Land-based sources of marine pollution: Pesticides, PAHs and phthalates in coastal stream water, and heavy metals in coastal stream sediments in American Samoa. Marine Pollution Bulletin, 116(1–2), 501–507. 10.1016/j.marpolbul.2016.12.058

Ponce-Vélez, G., & de la Lanza-Espino, G. (2019). Organophosphate Pesticides in Coastal Lagoon of the Gulf of Mexico. Journal of Environmental Protection, 10(02), 103–117. 10.4236/jep.2019.102007

Posit team (2023). RStudio: Integrated Development Environment for R. Posit Software, PBC, Boston, MA. URL http://www.posit.co/

Prakash, Sadguru. "Toxic effect of chlorpyrifos pesticides on the behaviour and serum biochemistry of Heteropnetues fossilis (Bloch)." International journal on agricultural Sciences 11, no. 1 (2020): 22–27.

Qiao, Kun, Tiantian Hu, Yao Jiang, Jianping Huang, Jingjin Hu, Wenjun Gui, Qingfu Ye, Shuying Li, and Guonian Zhu. "Crosstalk of cholinergic pathway on thyroid disrupting effects of the insecticide chlorpyrifos in zebrafish (Danio rerio)." Science of The Total Environment 757 (2021): 143769.

Reiss, Richard, Barbara Neal, James C. Lamb IV, and Daland R. Juberg. "Acetylcholinesterase inhibition dose–response modeling for chlorpyrifos and chlorpyrifos-oxon." Regulatory Toxicology and Pharmacology 63, no. 1 (2012): 124–131.

Robinson, Mark D., and Alicia Oshlack. "A scaling normalization method for differential expression analysis of RNA-seq data." Genome biology 11 (2010): 1–9.

Robinson, Mark D., Davis J. McCarthy, and Gordon K. Smyth. "edgeR: a Bioconductor package for differential expression analysis of digital gene expression data." bioinformatics 26, no. 1 (2010): 139–140.

Roche, H., Salvat, B., & Ramade, F. (2011). Assessment of the pesticides pollution of coral reefs communities from french polynesia. Revue d’Ecologie (La Terre et La Vie), 66(1), 3–10. http://documents.irevues.inist.fr/handle/2042/55860

Rohlf, F.J. and Marcus, L.F., 1993. A revolution morphometrics. Trends in ecology & evolution, 8(4), pp.129–132.

Roux, Natacha, Saori Miura, Mélanie Dussenne, Yuki Tara, Shu-hua Lee, Simon de Bernard, Mathieu Reynaud et al. "The multi-level regulation of clownfish metamorphosis by thyroid hormones." Cell reports 42, no. 7 (2023).

Roux, Natacha, Valentin Logeux, Nancy Trouillard, Rémi Pillot, Kévin Magré, Pauline Salis, David Lecchini, Laurence Besseau, Vincent Laudet, and Pascal Romans. "A star is born again: Methods for larval rearing of an emerging model organism, the False clownfish Amphiprion ocellaris." Journal of Experimental Zoology Part B: Molecular and Developmental Evolution 336, no. 4 (2021): 376–385.

Roux, N., Salis, P., Lambert, A., Logeux, V., Soulat, O., Romans, P., Frédérich, B., Lecchini, D., & Laudet, V. (2019). Staging and normal table of postembryonic development of the clownfish (Amphiprion ocellaris). Developmental Dynamics, 248(7), 545–568. 10.1002/DVDY.46

Rudis, Bob, Ben Bolker, Jan Schulz, A. Kothari, and J. Sidi. "ggalt: extra coordinate systems, geoms’, statistical transformations, scales and fonts for ‘ggplot2’." Available at: <https://CRAN.R-project.org/package=ggalt> (2017).

Ryu, Taewoo, Marcela Herrera, Billy Moore, Michael Izumiyama, Erina Kawai, Vincent Laudet, and Timothy Ravasi. "A chromosome-scale genome assembly of the false clownfish, Amphiprion ocellaris." G3 12, no. 5 (2022): jkac074.

Sabdono, A., Radjasa, O. K., Trianto, A., Sarjito, Munasik, & Wijayanti, D. P. (2019). Preliminary study of the effect of nutrient enrichment, released by marine floating cages, on the coral disease outbreak in Karimunjawa, Indonesia. Regional Studies in Marine Science, 30, 100704. 10.1016/J.RSMA.2019.100704

Salis, Pauline, Thibault Lorin, Victor Lewis, Carine Rey, Anna Marcionetti, Marie Line Escande, Natacha Roux et al. "Developmental and comparative transcriptomic identification of iridophore contribution to white barring in clownfish." Pigment Cell & Melanoma Research 32, no. 3 (2019): 391–402.

Salis, P., Roux, N., Soulat, O., Lecchini, D., Laudet, V., & Frédérich, B. (2018). Ontogenetic and phylogenetic simplification during white stripe evolution in clownfishes. BMC Biology, 16(1), 1–13. 10.1186/s12915-018-0559-7

Samajdar, Ishita, and Dipak Kumar Mandal. "Acute toxicity and impact of an organophosphate pesticide, chlorpyrifos on some haematological parameters of an Indian minor carp, Labeo bata (Hamilton 1822)." International journal of environmental sciences 6, no. 1 (2015): 106–113.

Sanden, M., P. A. Olsvik, L. Søfteland, J. D. Rasinger, G. Rosenlund, B. Garlito, Maria Ibáñez, and Marc HG Berntssen. "Dietary pesticide chlorpyrifos-methyl affects arachidonic acid metabolism including phospholipid remodeling in Atlantic salmon (Salmo salar L.)." Aquaculture 484 (2018): 1–12.

Saunders, L. M., Mishra, A. K., Aman, A. J., Lewis, V. M., Toomey, M. B., Packer, J. S., Qiu, X., McFaline-Figueroa, J. L., Corbo, J. C., Trapnell, C., & Parichy, D. M. (2019). Thyroid hormone regulates distinct paths to maturation in pigment cell lineages. ELife, 8. 10.7554/eLife.45181

Sayols, Sergi, Denise Scherzinger, and Holger Klein. "dupRadar: a Bioconductor package for the assessment of PCR artifacts in RNA-Seq data." BMC bioinformatics 17 (2016): 1–5.

Schindelin, Johannes, Ignacio Arganda-Carreras, Erwin Frise, Verena Kaynig, Mark Longair, Tobias Pietzsch, Stephan Preibisch et al. "Fiji: an open-source platform for biological-image analysis." Nature methods 9, no. 7 (2012): 676–682.

Schunter, Celia, Jennifer M. Donelson, Philip L. Munday, and Timothy Ravasi. "Resilience and Adaptation to Local and Global Environmental Change." In Evolution, Development and Ecology of Anemonefishes, pp. 253–274. CRC Press, 2022.

Shaw, M., Furnas, M. J., Fabricius, K., Haynes, D., Carter, S., Eaglesham, G., & Mueller, J. F. (2010). Monitoring pesticides in the Great Barrier Reef. Marine Pollution Bulletin, 60(1), 113–122. 10.1016/J.MARPOLBUL.2009.08.026

Sheikh, M.A., Fujimura, H., Miyagi, T., Uechi, Y., Yokota, T., Yasumura, S. and Oomori, T., 2009. Detection and ecological threats of PSII herbicide diuron on coral reefs around the Ryukyu Archipelago, Japan. Marine pollution bulletin, 58(12), pp.1922–1926.

Shu, Yan, Julin Yuan, Christer Hogstrand, Zhiyu Xue, Xilan Wang, Chunsheng Liu, Tao Li, Dapeng Li, and Liqin Yu. "Bioaccumulation and thyroid endcrione disruption of 2-ethylhexyl diphenyl phosphate at environmental concentration in zebrafish larvae." Aquatic Toxicology 267 (2024): 106815.

Soneson, C., Love, M. I., & Robinson, M. D. (2015). Differential analyses for RNA-seq: transcript-level estimates improve gene-level inferences. F1000Research, 4.

Soum, Thorn, Raymond James Ritchie, Raphatphorn Navakanitworakul, Sakshin Bunthawin, and Vipawee Dummee. "Acute toxicity of chlorpyrifos (CPF) to juvenile Nile tilapia (Oreochromis niloticus): genotoxicity and histological studies." (2022): 130–140.

Sudhakaran, Bindu Vijayakumari, and Sreeja S. "Impact of acute chlorpyrifos toxicity on the histology and biochemistry of the stomach, intestine, and muscle of the Anabas testudineus." bioRxiv (2023): 2023–10.

Sultatos, Lester G. "Metabolic activation of the organophosphorus insecticides chlorpyrifos and fenitrothion by perfused rat liver." Toxicology 68, no. 1 (1991): 1–9.

Tanvir, E. M., Rizwana Afroz, M. A. Z. Chowdhury, S. H. Gan, N. Karim, M. N. Islam, and M. I. Khalil. "A model of chlorpyrifos distribution and its biochemical effects on the liver and kidneys of rats." Human & experimental toxicology 35, no. 9 (2016): 991–1004.

Trasande, L., 2017. When enough data are not enough to enact policy: The failure to ban chlorpyrifos. PLoS Biology, 15(12), p.e2003671.

Triassi, M., Nardone, A., Giovinetti, M. C., de Rosa, E., Canzanella, S., Sarnacchiaro, P., & Montuori, P. (2019). Ecological risk and estimates of organophosphate pesticides loads into the Central Mediterranean Sea from Volturno River, the river of the “Land of Fires” area, southern Italy. Science of the Total Environment, 678, 741–754. 10.1016/j.scitotenv.2019.04.202

Tornero, V., & Hanke, G. (2016). Chemical contaminants entering the marine environment from sea-based sources: A review with a focus on European seas. In Marine Pollution Bulletin (Vol. 112, Issues 1–2, pp. 17–38). Pergamon. 10.1016/j.marpolbul.2016.06.091

Ubaid Ur Rahman, H., Asghar, W., Nazir, W., Sandhu, M. A., Ahmed, A., & Khalid, N. (2021). A comprehensive review on chlorpyrifos toxicity with special reference to endocrine disruption: Evidence of mechanisms, exposures and mitigation strategies. The Science of the Total Environment, 755(Pt 2), 142649. 10.1016/j.scitotenv.2020.142649

Velmurugan, Babu, Elif İpek Satar, Murat Yolcu, and Ersin Uysal. "Toxic impact of chlorpyrifos, an organophosphate pesticide, on serum biochemical parameters of Anabas testudineus." Acta Biologica Turcica 35, no. 1 (2022): 5–13.

Wang, Ruike, Mengxue Yang, Ye Zheng, Fuyong Song, Xiulan Zhao, and Chen Chen. "Interactive transgenerational effects of parental co-exposure to prochloraz and chlorpyrifos: Disruption in multiple biological processes and induction of genotoxicity." Pesticide Biochemistry and Physiology 198 (2024): 105713.

Wang, Yao, Shaochun Yuan, Xin Jia, Yong Ge, Tao Ling, Meng Nie, Xihong Lan, Shangwu Chen, and Anlong Xu. "Mitochondria-localised ZNFX1 functions as a dsRNA sensor to initiate antiviral responses through MAVS." Nature cell biology 21, no. 11 (2019): 1346–1356.

Wickham, Hadley. Data analysis. Springer International Publishing, 2016.

Wingett, Steven W., and Simon Andrews. "FastQ Screen: A tool for multi-genome mapping and quality control." F1000Research 7 (2018).

Wołejko, E., Łozowicka, B., Jabłońska-Trypuć, A., Pietruszyńska, M., & Wydro, U. (2022). Chlorpyrifos Occurrence and Toxicological Risk Assessment: A Review. In International Journal of Environmental Research and Public Health (Vol. 19, Issue 19, p. 12209). Multidisciplinary Digital Publishing Institute. 10.3390/ijerph191912209

Wong, H. L., Garthwaite, D. G., Ramwell, C. T., & Brown, C. D. (2018). Assessment of occupational exposure to pesticide mixtures with endocrine-disrupting activity. Environmental Science and Pollution Research 2018 26:2, 26(2), 1642–1653. 10.1007/S11356-018-3676-5

Wu, Tianzhi, Erqiang Hu, Shuangbin Xu, Meijun Chen, Pingfan Guo, Zehan Dai, Tingze Feng et al. "clusterProfiler 4.0: A universal enrichment tool for interpreting omics data." The innovation 2, no. 3 (2021).

Xiong, Jingyuan, Liantian Tian, Yongjie Qiu, Ding Sun, Hao Zhang, Mei Wu, and Jintao Wang. "Evaluation on the thyroid disrupting mechanism of malathion in Fischer rat thyroid follicular cell line FRTL-5." Drug and Chemical Toxicology 41, no. 4 (2018): 501–508.

Xu, J., Ke, Z., Xia, J., He, F., & Bao, B. (2016). Change of body height is regulated by thyroid hormone during metamorphosis in flatfishes and zebrafish. General and Comparative Endocrinology, 236, 9–16. 10.1016/j.ygcen.2016.06.028

Yin, XiaoHui, GuoNian Zhu, Xian Bing Li, and ShaoYing Liu. "Genotoxicity evaluation of chlorpyrifos to amphibian Chinese toad (Amphibian: Anura) by comet assay and micronucleus test." Mutation Research/Genetic Toxicology and Environmental Mutagenesis 680, no. 1-2 (2009): 2–6.

Yu, Guangchuang, Li-Gen Wang, Yanyan Han, and Qing-Yu He. "clusterProfiler: an R package for comparing biological themes among gene clusters." Omics: a journal of integrative biology 16, no. 5 (2012): 284–287.

Zwahlen, J., Gairin, E., Vianello, S., Mercader, M., Roux, N. and Laudet, V., 2024. The ecological function of thyroid hormones. Philosophical Transactions of the Royal Society B, 379(1898), p.20220511.

