## Supplemental information for "The multi-level effect of chlorpyrifos during clownfish metamorphosis"

**Supporting Information files: The multi-level effect of chlorpyrifos during clownfish metamorphosis (Reynaud-Vianelloetal)**

**Supplementary Materials and Methods**

1. Animal Rearing methods

1.1. Licences/Ethics Approval

*A.ocellaris* reproductive couples and larvae were reared under licence number 21-12-1769 issued from Academia Sinica’s Institutional Animal Care and Use Committee to the Marine Eco-Evo-Devo Unit at the LinHai Marine Research Station; or from approval number A6601601 issued from the C2EA-36 Ethics Committee for Animal Experiment Languedoc-Roussillon (CEEA-LR). All experiments were performed within the provisions of the same licences.

1.2. Details of animals used and pharmacological treatments

*A.ocellaris* reproductive couples were bought from a commercial anemonefish farm (S.T Biotechnology Inc; Changhua County, Taiwan). The reproductive couple was housed in a 38L tank in recirculating natural seawater under a 15h:9h day-night photoperiod, at 28°C. Fish were fed thrice daily either with dry pellets (Hikari® Premium Megabite; Kyorin Food Industries, Ltd) or with fresh seafood (homemade). At spawning fish laid eggs on a terracotta pot present in the tank, which was then transferred to a dedicated hatching tank 7-8 days later (based on observed progression of embryonic development). Larvae hatched in a temperature-controlled and oxygenated tank, shielded from light, in natural sea water supplemented daily with vitamin B12- and DHA/EPA-enriched unicellular green algae *Chlorella spp.* (Super-Fresh Chlorella V12 Solution; Chlorella Industry Co. Ltd) and live S-type rotifers *Brachionus rotundiformis* (grown in house), themselves enriched with instant dry yeast. In either facility, larvae were collected at 5dph (white bar experiment) or 8dph (RNAseq experiment) and randomly transferred to dedicated beakers for pharmacological treatment.

Pharmacological treatment was performed in a small-volume setup analogous to what described in Roux et al., 2021. Briefly, larvae were maintained and reared in separate, oxygenated 1L glass beakers (around 30 larvae per condition, no more than 15 larvae per beaker) in a shared tank partially filled with temperature-controlled seawater (thus maintaining constant temperature within the beakers by conduction). Water that evaporated from the beakers over time was replaced daily with the correct volume of seawater-chemical compound solution. Larvae in beakers were fed daily with live rotifers, and — from 8dph onwards — with live Artemia franciscana nauplii (freshly hatched from dehydrated eggs; Golden West Artemia Supreme Plus/Great Salt Lake Artemia). Caution was taken to maintain high beaker water quality.

2. RNA extraction and sequencing details

2.1 RNA extraction

On the day of extraction, sample lysates (in Trizol) were thawed on ice and centrifuged 10 min in a tabletop microcentrifuge at 11000 x g, 4°C; as to collect any leftover tissue debris to the bottom. 750µL of each supernatant were loaded into a dedicated RNA filtering column (NucleoSpin® RNA Mini kit; Macherey-Nagel CAT#740955.50), and processed according to manufacturer recommendations. Specifically — for each sample — 350uL of 70% Ethanol (Honeywell/Riedel-deHaen CAT#32221, in ddH2O) were added to the filtered flowthrough, the solution was then vortexed, and loaded onto a RNA binding column. After desalting (kit-supplied Membrane Desalting Buffer), contaminating DNA was digested by a 15 min incubation in recombinant DNAse (in Reaction Buffer, room temperature). Guanidine hydrochloride/ethanol wash buffers (RAW2 and then, twice, RA3) were then sequentially used to denature proteins, discarding the flowthrough from centrifugation after each step. After a further centrifugation to remove all possible leftover buffer/ethanol (2 min, 11000 x g, 4°C), membrane-bound RNA was eluted in 30uL of RNAse-free water (UltraPure™ DNase/RNase-Free Distilled Water; Invitrogen™/Thermo Fisher Scientific CAT#10977015), in a 1.5mL RNAse-free tube (kit-supplied). RNA amounts and quality were initially assessed with a NanoDrop spectrophotometer (NanoDrop Lite; Thermo Scientific™ CAT#840281500) and by agarose gel electrophoresis (2:1 intensity ratio of 28S:18S bands; 800ng of sample per lane, 0.6 % Agarose in TAE-buffer, 1h, 50V). Extracted RNA was stored at -80C until sequencing.

2.2 Library preparation and sequencing

Quality-control, library preparation and sequencing of extracted RNA were performed by the High Throughput Genomics Core at Academia Sinica, Taipei. RNA-Seq libraries were generated from 2000 ng of total RNA using the Illumina Stranded mRNA Prep mRNA Sample Preparation Kit with UDI indices (Illumina, USA) according to manufacturer's instructions. Surplus PCR primers were removed using AMPure XP Bead-Based Reagent (Beckman Coulter Life Sciences, USA). Final cDNA libraries were checked for quality and quantified using Qubit (ThermoFisher Scientific, USA) and Fragment Analyzer for size profiling (Agilent, USA), and concentration-normalised using KAPA Library Quantification Kit for Illumina Platforms (Roche, USA). Sequencing was performed on an Illumina NextSeq2000 for paired-end 150 base format. Libraries were loaded in a P3 flow cell at 644 pM. The fastQ files were generated and demultiplexed using the Illumina bcl2fastq v2.20 pipeline. A median of 24 million paired-end reads per sample were obtained (expected output ceiling: 28 million reads/sample).

2.3. Analysis of bulkRNAseq data

The complete Rnotebooks use for the analysis of the data (also indicating the versions of all packages used), the counts matrix, the gene and sample metadata files, as well as other data to reproduce the analysis are available at [https://github.com/StefanoVianello/ReynaudVianello_AoceCPF](about:blank); the pipeline is summarised below.

2.4. Pre-processing

Raw (demultiplexed) fastq files were quality-checked based on reports generated by using FastQC v0.12.0 (RRID:SCR_014583; Andrews, 2010) with default parameters, before and after adapter trimming. Adapters were trimmed using the function bbduk (RRID:SCR_016969) of BBTools v39.01 (RRID:SCR_016968; Bushnell B., [http://sourceforge.net/projects/bbmap/](about:blank)) with ktrim=r, and k=23. The flag trimpolyg=40 was also used, given that the initial FastQC run had identified over-representation of polyG sequences (in fact missing data, due to 2 defectuous tiles of the patterned flow cell). Yield and quality of the run were well within Illumina’s specification ( >1000M PF fragments and Q30 bases >75%). Categories flagged by FastQC after trimming (“warning” or “fail”) were analysed in detail and judged not to be prejudicial to further analysis (see dedicated Rnotebook). Issues relating to the detection of high sequence duplication rates were diagnosed based on the output of the analyzeDuprates function of the dupRadar package (Sayols et al., 2016). To this aim, duplicate reads were marked with the function markdup from SAMtools v1.18 (RRID:SCR_002105; Danecek et al; 2021) with default parameters.

As a further quality control step to detect sample contamination, trimmed reads were also run through FastQ Screen v0.15.2 (RRID:SCR_000141; Wingett, Andrews, 2018) with default parameters, against a manually curated set of genomes including — in addition to default ones — those of other fish species used by neighbouring laboratories (goldfish *Carassius auratus*: ASM336829v1, carp *Cyprinus carpio*: ASM1834038v1, zebrafish *Danio rerio*: GRCz11), the three main live foods fed to our fish (*Artemia franciscana*: AFR02, *Brachionus rotundiformis*: ASM1680229v1, *Chlorella vulgaris*: cvul), and additional possible contaminating species (ant *Formica exsecta*: ASM365146v1). All FastQ Screen genomes were indexed with bowtie2 v2.4.4 (RRID:SCR_016368; Langmead, Salzberg, 2012) as per FastQ Screen recommendation.

**Supplementary Tables**

| *Predictors* | *Df* | *SS* | *MS* | *Rsq* | *F* | *Z* | *P-value* |
| --- | --- | --- | --- | --- | --- | --- | --- |
| *Time* | *1* | *0.176124* | *0.176124* | *0.286333* | *129.5786* | *5.028655* | *0.001* |
| *Treatments* | *3* | *0.042403* | *0.014134* | *0.068936* | *10.39888* | *8.407942* | *0.001* |
| *Time:Treatments* | *3* | *0.009201* | *0.003067* | *0.014958* | *2.256443* | *3.452869* | *0.001* |
| *Residuals* | *285* | *0.387373* | *0.001359* | *0.629772* | */* | */* | */* |
| *Total* | *292* | *0.6151* | */* | */* | */* | */* | */* |

***Table S1: Treatment and time effect on shape variation during A. ocellaris metamorphosis, assessed by multivariate models.***

**Supplementary figures**

**Figure S1**

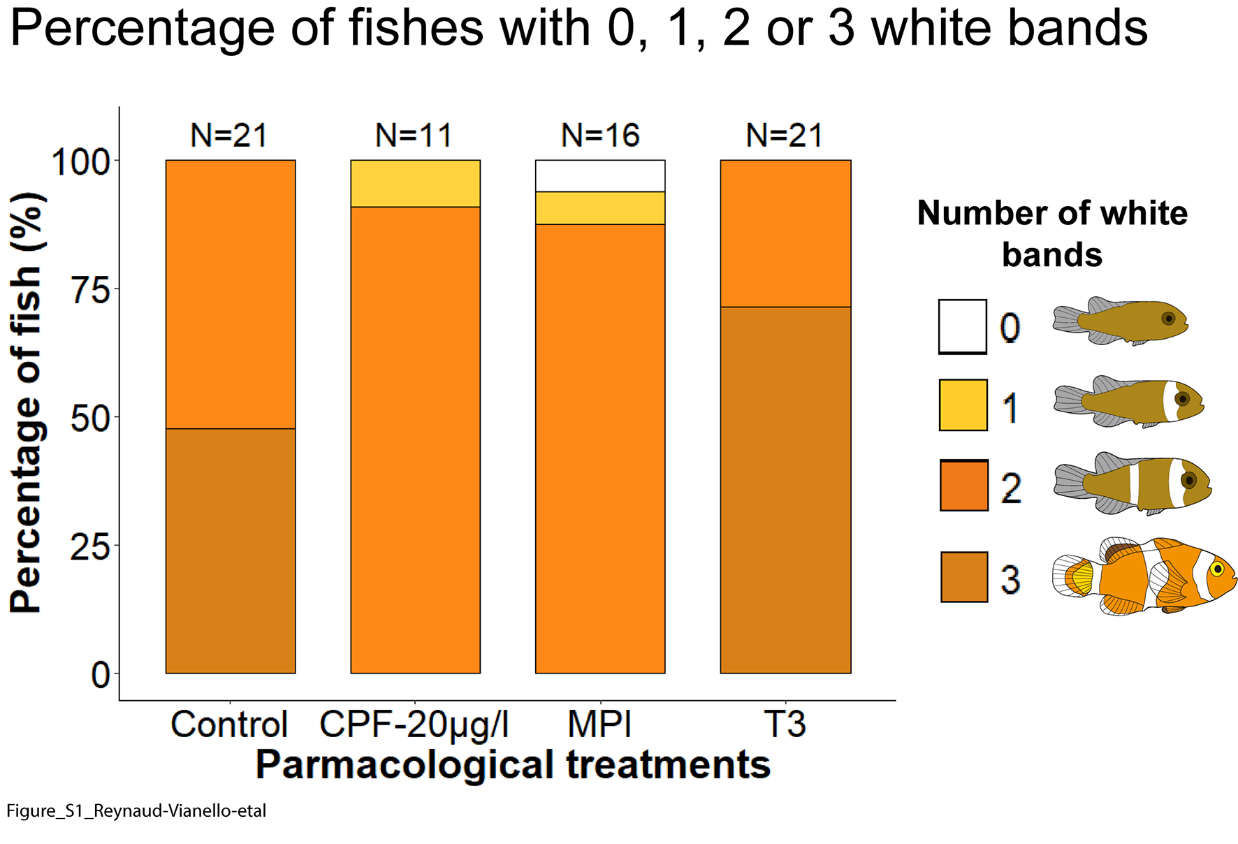

**Figure S2**

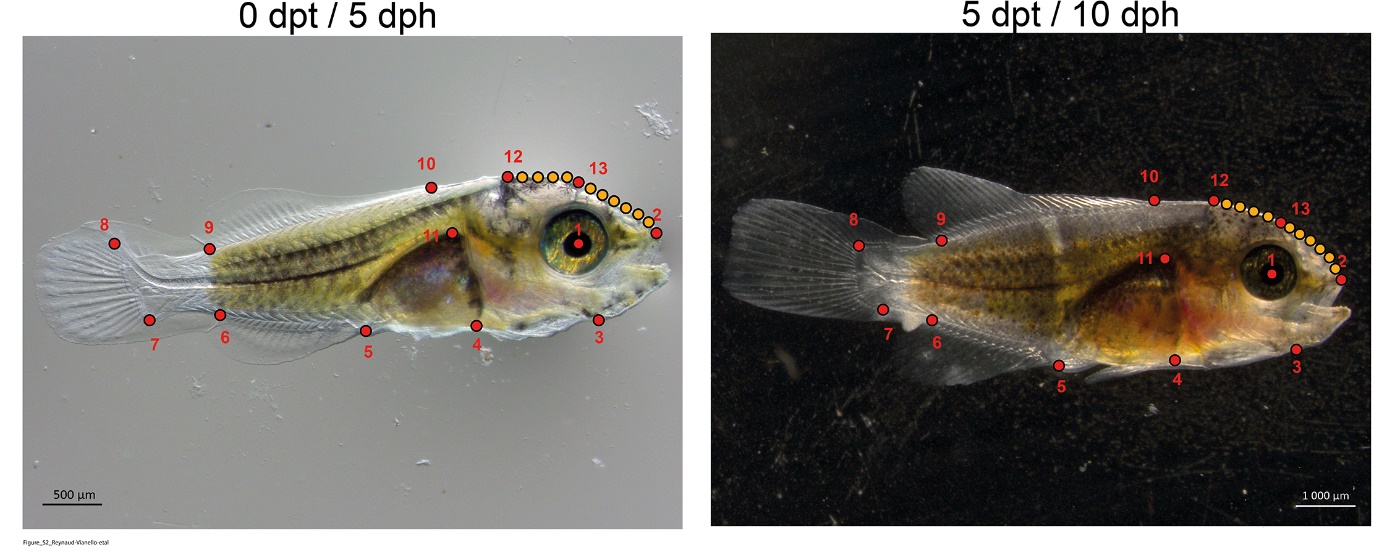

**Figure S3**

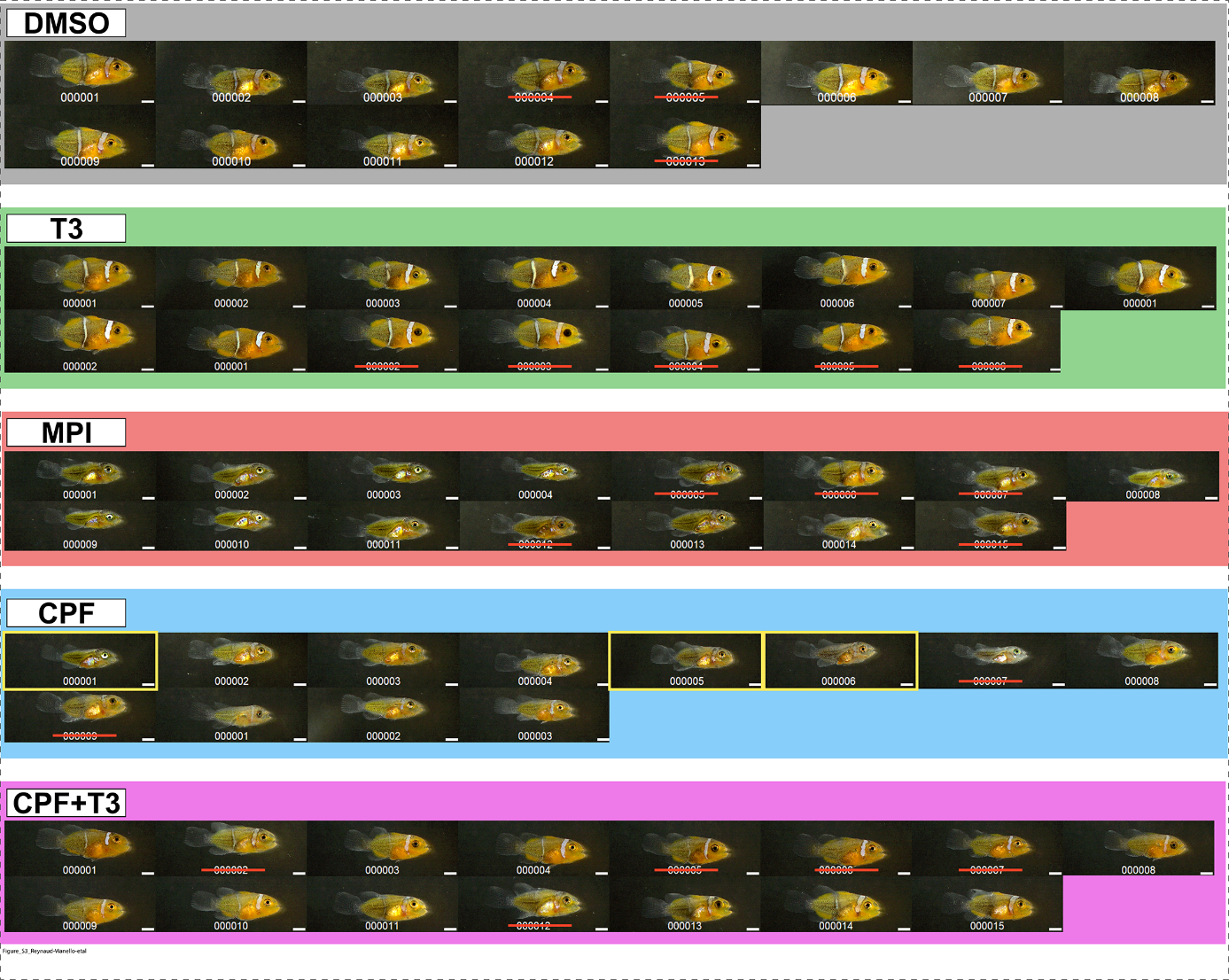

**Figure S4**

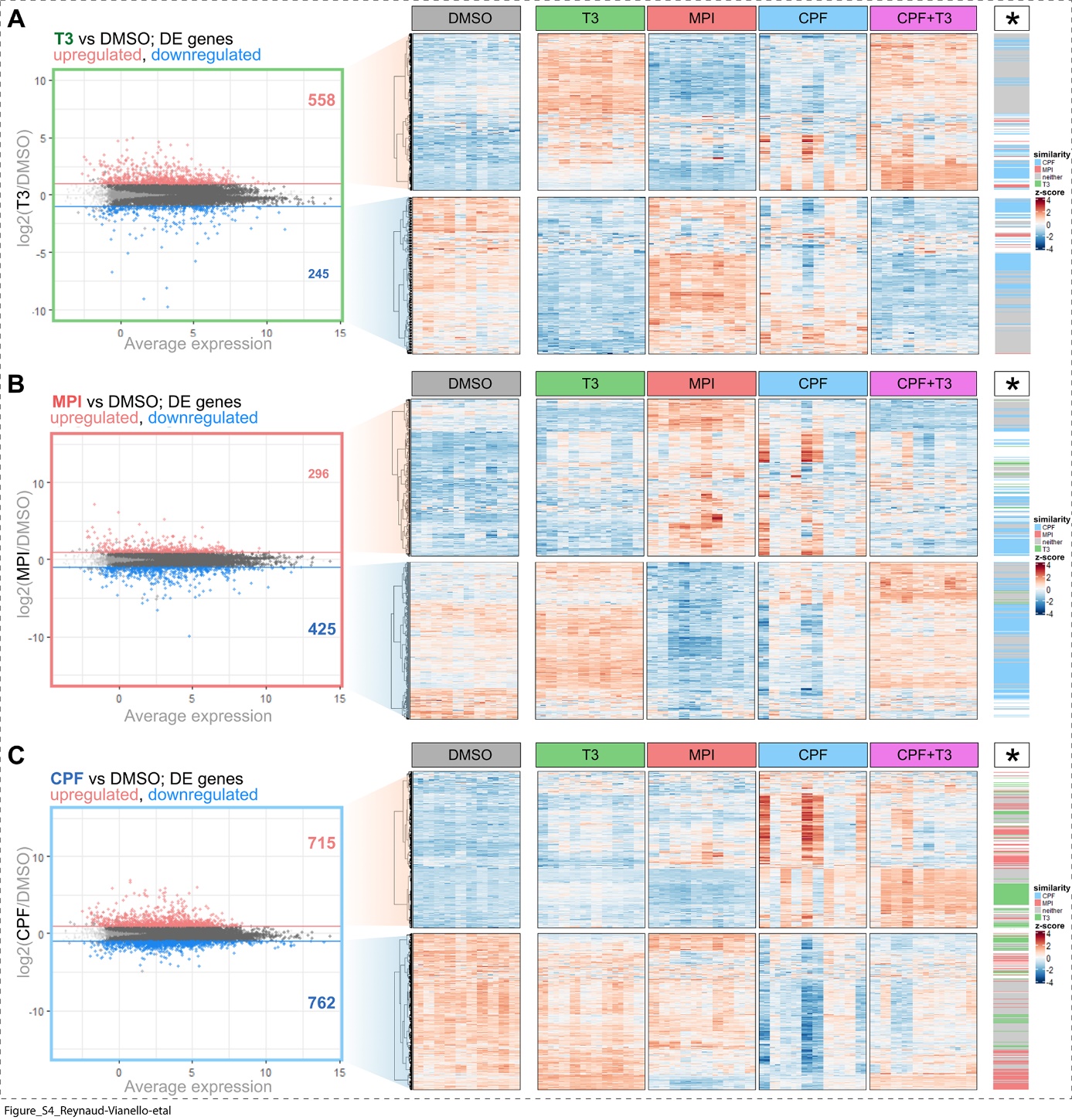
